# Non-coding RNA RsaE regulates biofilm thickness, viability and dissemination in methicillin-resistant *Staphylococcus aureus*

**DOI:** 10.64898/2026.03.05.709773

**Authors:** Mehak Chauhan, Ivayla Ivanova, Emily G. Sudnick, Ryan W. Steere, Julia R. Tennant, Jacob A. Hensley, Pedro Arede, Gregory M. Jensen, Isabelle Hatin, Olivier Namy, Philippe Bouloc, Ronan K. Carroll, Sander Granneman

## Abstract

Methicillin-resistant *Staphylococcus aureus* (MRSA) is a formidable human pathogen responsible for life-threatening infections worldwide. Central to its pathogenic success is the tightly coordinated regulation of virulence factors, including the phenol soluble modulins (PSM), short amphipathic toxins that drive cytolysis, immune evasion and biofilm maturation. We previously identified the conserved non-coding RNA RsaE as a putative regulator of the *psm⍺* operon, but the biological significance of this interaction remained unclear. Here we show that RsaE and endoribonuclease Y jointly regulate *psm⍺* transcript abundance at the post-transcriptional and transcriptional levels, the latter primarily through activation of the *agr* quorum sensing system. We show that the *psm⍺* transcript is unusually stable and highly structured, with Shine-Dalgarno sequences of individual toxin-coding sequences differentially accessible, providing a mechanism for translational fine-tuning of individual PSM⍺ peptides. *In vitro* biofilm analyses revealed that RsaE deletion produces thinner biofilms with reduced extracellular DNA accumulation on the surface and transiently elevated cell viability. Strikingly, in a murine catheter infection model, *ΔrsaE* biofilms exhibited structural abnormalities and significantly reduced dissemination to kidneys. These findings identify the RsaE non-coding RNA as a key regulator linking central metabolism and quorum sensing to toxin expression, biofilm maturation and infection in MRSA.

**Author summary:** Methicillin-resistant *Staphylococcus aureus* (MRSA) causes severe, often device-associated infections that are hard to treat. A central reason for its success is tight control over toxins such as the ⍺-phenol-soluble modulins (PSM⍺), which help bacteria damage host cells and build/disperse biofilms. Small RNAs (sRNAs) are fast-acting genetic regulators in bacteria, that also regulate toxin production. We show that the conserved sRNA RsaE and the endoribonuclease RNase Y jointly tune the level and lifetime of the *psm⍺* RNA transcript, which encodes four PSM⍺ peptides. The *psm⍺* RNA is unusually stable and highly structured with its design favouring translation of PSM⍺4 over the other peptides. Removing RsaE or RNase Y increases *psm⍺* transcription via the *agr* quorum-sensing system and further stabilises the *psm⍺* RNA. In biofilms, loss of RsaE reduces the thickness and extracellular DNA layer, while increasing early cell viability. In a mouse catheter model, RsaE deletion leads to structurally altered biofilms and reduced early dissemination to kidneys.

These results identify RsaE as a key regulator connecting metabolism and quorum sensing to toxin expression, biofilm maturation and disease progression, and point to RNA-centred strategies to regulate MRSA spread from biofilms.

## Introduction

*Staphylococcus aureus*, an opportunistic Gram-positive bacterium is responsible for a wide array of human diseases ranging from minor skin infections to life-threatening pneumonia. It is also the most common cause of secondary infections in COVID-19 patients [1]. Its resistance to multiple and diverse drugs continues to be a significant healthcare burden globally, yet few new antimicrobials have been developed to battle MRSA infections [2]. This diverse range of manifested diseases is due to the production of many virulence factors as well as the coordinated expression of multiple regulators [3].

Small regulatory RNAs (sRNAs) play highly significant roles in regulating bacterial gene regulation and adaptation to diverse environments [4]. In *S. aureus*, they act as a fast regulatory mechanism that helps the bacterium sense and respond to rapid environmental changes such as nutrient limitation, oxidative stress, and the host immune response [5] They do so largely by influencing mRNA stability and translation [6–8]. One of the best studied examples is the sRNA RNAIII, the principal effector of the accessory gene regulator (*agr*) quorum-sensing system. It encodes a toxin itself (*hld*) but as a regulatory RNA also coordinates the expression of many other toxins and surface proteins critical for virulence [9]. Comparative and transcriptomic studies have revealed many other sRNAs, termed RsaA-K, each with distinct regulatory functions [10,11]. Among them, RsaE has drawn particular interest because of its conserved function in Gram-positive bacteria and its role in regulating many metabolic pathways involving folate metabolism, the TCA cycle [10,12,13]. RNA-interactome analyses, using UV cross-linking, ligation and sequencing of hybrids (CLASH), showed that RsaE also base-pairs with *psm⍺* operon transcripts, that encode cytolytic phenol-soluble modulins (PSMs) [14]. PSM⍺ 1-4 are short, amphipathic ⍺-helical peptides that play distinct and diverse roles in *S. aureus* [15]. They act as cytolytic toxins capable of lysing erythrocytes and neutrophils and help to structure and mature biofilms as well as dissemination from biofilms [16–18]. Even though their structure and physiochemical properties are very similar, they vary in the functions they perform. Whereas ⍺2 and ⍺3 peptides are the most cytolytic in nature, ⍺3 forms ⍺ amyloid fibrils whereas ⍺1 and ⍺4 form β fibrils, which provide stability to the biofilms [15,19,20].

Because PSM⍺1-4 are encoded on a single operon, it was assumed that they are expressed at similar levels. However, PSM⍺2 and ⍺3 are less abundantly present in culture supernatants compared to PSM⍺1 and 4 [14,21,22], hinting that they are differentially expressed and/or secreted. In the community acquired USA300 genetic background (CA-MRSA), deleting RsaE reduces the accumulation of PSM⍺1 and 4 in culture supernatants, suggesting a regulatory role for RsaE in modulating the production of individual PSM⍺ peptide levels [14]. However, the functional significance of this observation remained unresolved. Given that these toxins are secreted at high levels can be harmful to the bacterium and each one has a distinct and diverse important function, we hypothesised that their expression must be carefully controlled. We hypothesised that RsaE acts as a molecular tuner of PSM⍺ expression, ensuring that these peptides are produced at optimal levels for MRSA adaptation to host environments and for the structuring and maturation of biofilms. To address this hypothesis, we investigated how RsaE regulates the *psm⍺* operon and examined the consequences of RsaE deletion on biofilm development both *in vitro* and *in vivo*. Collectively, our results reveal important roles for RsaE in regulating biofilm maturation, cell viability, and bacterial dissemination during infection.

## Results

### Mapping the *psm⍺* transcript boundaries

Before assessing the regulatory impact of RsaE on *psm⍺*, we characterised the *psm⍺* operon transcript in more detail. While the coding sequences for USA300 *psm⍺* peptides were annotated in USA300 genome files (USA300_FPR3757; https://aureowiki.med.uni-greifswald.de/SAUSA300_0424.4), the precise boundaries of the operon transcript were undefined. Previous work [23] also indicated that the USA300 *psm⍺* gene may contain an internal promoter that produces Sau-41, an sRNA predicted to be ∼150 nucleotide in length that initiates just upstream of *psm⍺4* (Fig 1a). However, USA300 transcription start site (TSS) mapping data [24] did not provide evidence for an internal TSS in this region, although a TSS was predicted upstream of *psm⍺2*. To resolve these conflicting observations, we undertook a detailed analysis of the *psm⍺* transcript.

**Fig 1.**
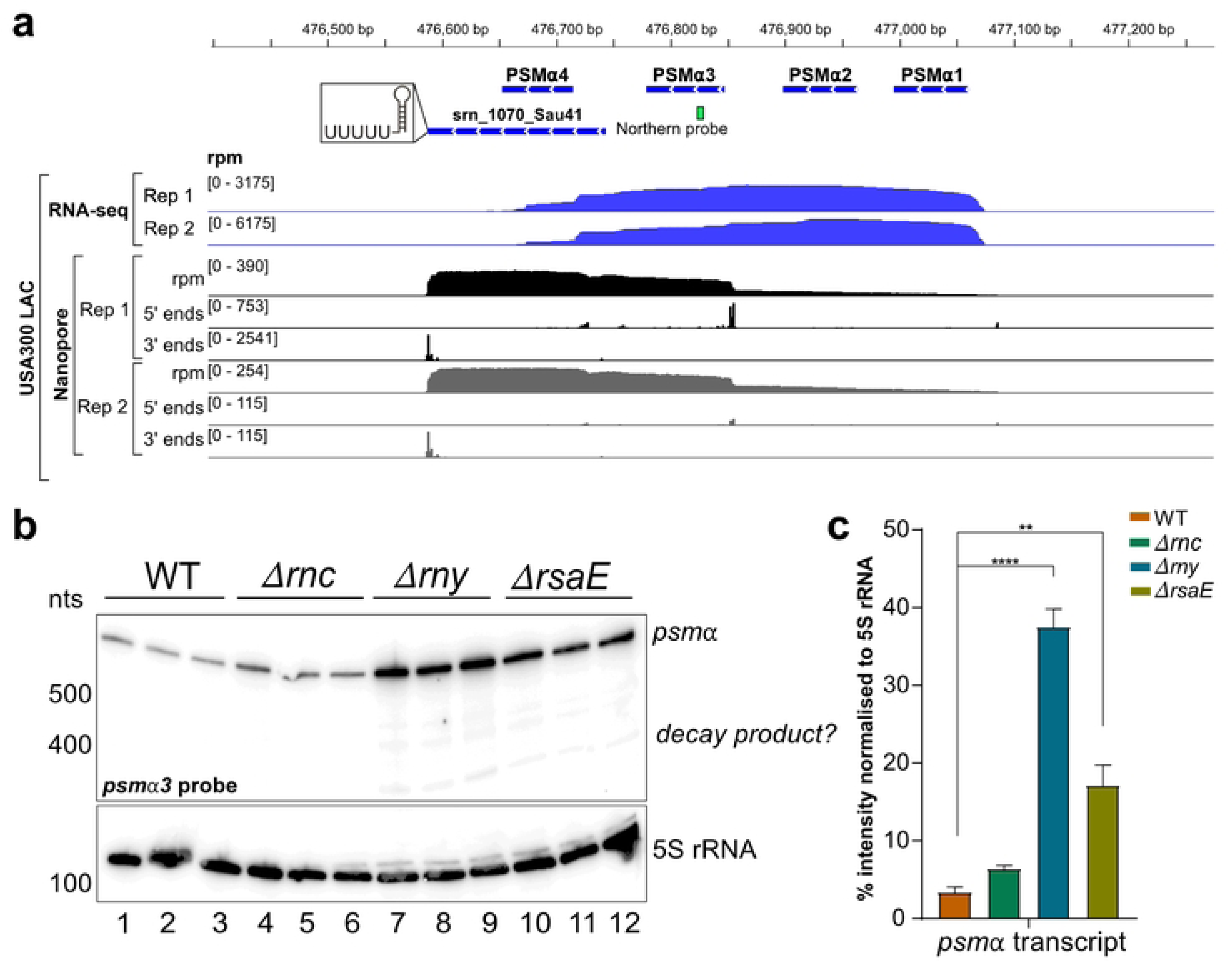
RNase Y and RsaE regulate *psm⍺* steady state RNA levels. **(a)** Genome browser visualisation of *psm⍺* transcript in wild-type. Shown is results from published RNA-seq and Nanopore data [14] from USA300 cells grown in TSB to OD_595_ ∼ 3.0. Read abundance is shown in reads per million (rpm). **(b)** Northern blot analysis of wild-type (WT), *ΔrsaE*, *Δrny* and *Δrnc* mutants. Three independent biological replicates were grown till late exponential phase (OD_595_ ∼ 3.0) in TSB and probed for *psm⍺3* transcript. **(c)** Quantification of Northern blot signals for *psm⍺3*. Bar plots show average percentage of signal intensities normalised to 5S rRNA, with standard deviations calculated from three independent biological replicates. P-values above the bars represent statistical comparisons between WT and mutants, obtained using two-way ANOVA (p < 0.01 = **, p < 0.0001 = ****).

To map the transcript boundaries and identify potential shorter isoforms, we first reanalysed our published RNA-seq and Nanopore datasets from USA300 grown in TSB medium [14]. To complement these analyses, we also performed Northern blot analyses using four different probes that hybridise to individual *psm⍺* coding sequences (Fig 1b, S1 Fig). While the short-read Illumina data showed more reads near the predicted 5’ end, the long-read Nanopore data better resolved the 3’ end with most reads terminating at the predicted transcription terminator (Fig 1a). Analysis of the cumulative counts of 5’ and 3’ ends of the Illumina and Nanopore reads indicated that the *psm⍺* transcript is approximately 514 nucleotides long. The Northern blot analyses showed that the most abundant *psm⍺* transcript is longer than 500 nucleotides, which presumably represents the ∼514 nt full-length transcript (Fig 1b, S1 Fig). Long-read data also revealed shorter *psm⍺* transcript fragments (∼380 nucleotides), with 5’ ends mapping to the SD sequence of *psm⍺3*. Northern blotting also revealed shorter *psm⍺* fragments (∼350-400 nt), but these were low abundant and could be detected with all probes that hybridise to individual *psm⍺* coding sequences (Fig 1b, S1a Fig). Under the tested growth conditions, we did not detect the reported ∼150 nucleotide Sau-41 sRNA (S1 Fig).

### RsaE and RNase Y regulate *psm⍺* at the transcriptional and post-transcriptional level

RNA-RNA interactome (CLASH; [14]) suggested that *S. aureus* RsaE base-pairs with the Shine-Dalgarno (SD) sequences of *psm⍺2* and *psm⍺3*. Furthermore, IntaRNA predictions indicate that RsaE base-pairs with the SD sequences of *psm⍺1-3* (S2 Fig). This implies that RsaE may regulate the stability and/or translation of *psm⍺*.

To investigate the impact of RsaE on *psm⍺* stability, we compared the *psm⍺* RNA steady state levels in USA300 and its *ΔrsaE* derivative. We also included *Δrnc* (RNase III) and *Δrny* (RNase Y) deletion mutants, as both endonucleases are key contributors to virulence by regulating the stability of numerous mRNAs. Moreover, CLASH data implied that RNase III mediates the RsaE-*psm⍺* interaction [14].

Deletion of *rsaE* significantly increased *psm⍺* steady state levels (Fig 1b, lanes 10-12 vs. 1-3; Fig 1c). In contrast, the *Δrnc* mutation had no significant impact on *psm⍺* steady-state levels (Fig 1b, lanes 4-6 vs. 1-3; Fig 1c). Strikingly, the *Δrny* strain exhibited a ∼10-fold increase in *psm⍺* expression compared to the parental (WT) strain (Fig 1b, lanes 7-9 vs. 1-3; Fig 1c).

Taken together, these results demonstrate that RsaE and RNase Y are negative regulators of *psm⍺* transcript levels.

### RsaE and RNase Y reduce the expression and half-life of the *psm⍺* transcript

The elevated *psm⍺* steady-state levels in the *Δrny* and *ΔrsaE* strains hinted that these genes play a role in degradation of the operon transcript. If this is correct, the half-life of the *psm⍺* transcript should increase significantly in their absence. To test this, we performed a rifampicin stability assay [25] to analyse the decay of the transcript. This antibiotic inhibits RNA polymerase, allowing us to monitor the rate of RNA decay over time. RNA was extracted at various time points up to one hour and *psm⍺* transcript levels were monitored by Northern blot analysis (Fig 2a).

**Fig 2.**
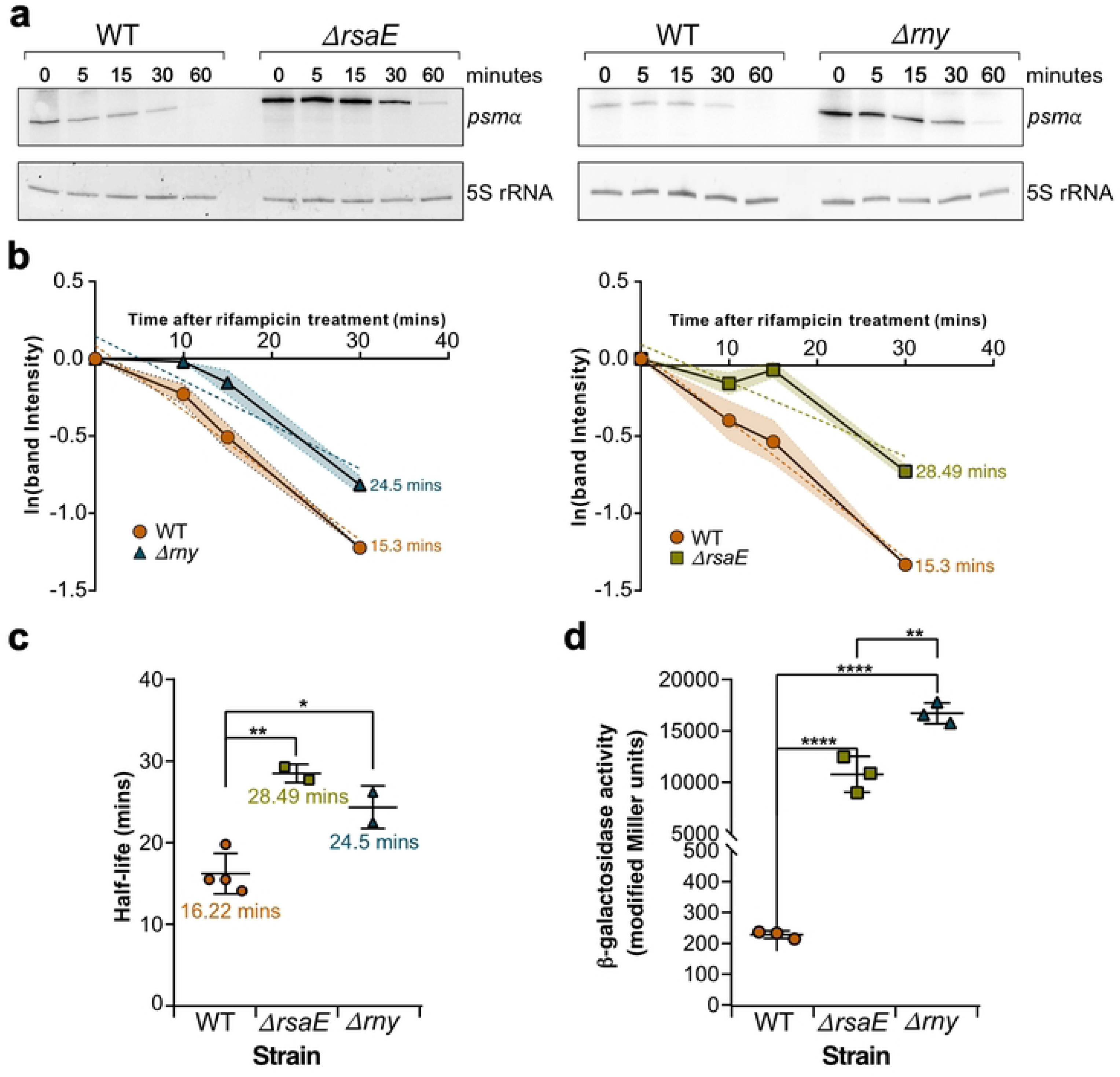
RsaE regulates *psm⍺* RNA transcription and half-life. **(a)** Rifampicin Stability assay. Total RNA from rifampicin-treated wild-type (WT), *ΔrsaE*, and *Δrny* strains (two biological replicates each) was collected at 0, 5, 15, 30, and 60-minutes posttreatment. RNA samples were fractionated on a 6% polyacrylamide-urea (PAA-urea) gel and probed with a radiolabelled *psm⍺* probe. 5S rRNA, visualised from a SYBR stained gel, was used as a loading control. **(b)** Natural log-transformed band intensity values were quantified using ImageJ. Decay slopes were calculated in GraphPad Prism, and transcript half-lives were derived from the slope of the linear regression. The shaded error region represents the standard deviation calculated from two biological replicates. **(c)** Bar plots showing the calculated half-lives of *psm⍺* transcripts in WT, *ΔrsaE*, and *Δrny* strains, based on two biological replicates per sample. Error bars represent standard deviations. Statistical significance was assessed using one-way ANOVA followed by post hoc Dunnett test (p < 0.01 = **, p < 0.001 = ***). **(d)** *psm⍺* promoter activity as represented by β-galactosidase activity in wild-type, *ΔrsaE*, and *Δrny* strains. Strains carried plasmid pJB185 with the *lacZ* gene under control of the *psm⍺* promoter. Enzyme activity was measured using three technical replicates from three biological replicates.

Since 90% of *S. aureus* RNAs have half-lives of less than 5 minutes [26], the persistence of the *psm⍺* transcript 30 minutes after drug treatment indicates exceptional stability. Quantification of the Northern blot data revealed that, under exponential growth in TSB, the *psm⍺* transcript has a half-life exceeding 15 minutes, assuming exponential decay (Fig 2b-c). In the *Δrny* and *ΔrsaE* strains, the *psm⍺* transcript exhibited biphasic decay kinetics: an initial stabilisation phase with minimal degradation during the first 15 minutes after rifampicin treatment, followed by accelerated decay. Fitting the complete decay curves to an exponential model revealed that the *psm⍺* transcript half-life was approximately twice as long in *Δrny* and *ΔrsaE* strains compared to the parental strain (∼25-28 minutes vs. ∼15 minutes; Fig 2b-c). When only considering the 0-, 5-and 15-minute time points, the *psm⍺* transcript remained substantially more stable in *Δrny* (apparent t_1/2_ = 80.53 minutes) and *ΔrsaE* (apparent t_1/2_ = 44.35 minutes) strains compared to the parental strain (t_1/2_ = 20.77 minutes). After the time point 15, the decay rates were similar across all strains.

To determine whether changes in *psm⍺* transcript steady-state levels were due solely to altered decay or also involved transcriptional changes, we directly measured *psm⍺* promoter activity using a transcriptional reporter construct, in which LacZ expression was driven by the *psm⍺* promoter. These analyses revealed significantly higher *psm⍺* promoter activity in *Δrny* and *ΔrsaE* strains (Fig 2d).

We conclude that RsaE and RNase Y regulate *psm⍺*transcript abundance at the post-transcriptional and transcriptional level.

### The *psm⍺* operon transcript is highly structured

The remarkably long half-life of the *psm⍺* transcript prompted us to study its structure in more detail. We hypothesised that extensive secondary structure formation contributes to its exceptional stability. Consistent with this idea, *in silico* analysis using RNAfold [27] predicted that a substantial fraction of the *psm⍺* transcript forms stable base-pairing interactions. Notably, all toxin SD sequences were predicted to be at least partially sequestered in secondary structures (S3a Fig). To experimentally validate this predicted structure, we performed *in vivo* SHAPE-MaP experiments using the USA300 parental strain and the 2-aminopyridine-3-carboxylic acid imidazolide (2A3) SHAPE reagent [28]. Sites of modification were mapped by Illumina high-throughput sequencing of the resulting cDNAs followed by mutational profiling analysis. The derived SHAPE reactivity values were then integrated into the RNA Framework package [29] as structural constraints to generate a refined *psm⍺* transcript secondary structure model.

The SHAPE-MaP-informed structure confirmed that the *psm⍺* transcript is highly structured, forming an elaborate network of stem-loops and helical regions (Fig 3a-b). Notably, all four start codons are sequestered within stem structures, with the SD sequences of *psm⍺2* and *psm⍺3* deeply embedded within stable base-paired stems, likely reducing their accessibility for ribosome binding. In contrast, the *psm⍺4* SD sequence resides in a flexible region, whereas the *psm⍺1* SD is predicted to only be partially sequestered in stem structures. This structural distinction was further supported SHAPE reactivity data: most nucleotides within the *psm⍺1-3* SD sequences showed low reactivity, indicating strong base-pairing, whereas three of six nucleotides in the *psm⍺4* SD showed higher reactivity, indicating greater structural flexibility (Fig 3b; S3b Fig).

**Fig 3.**
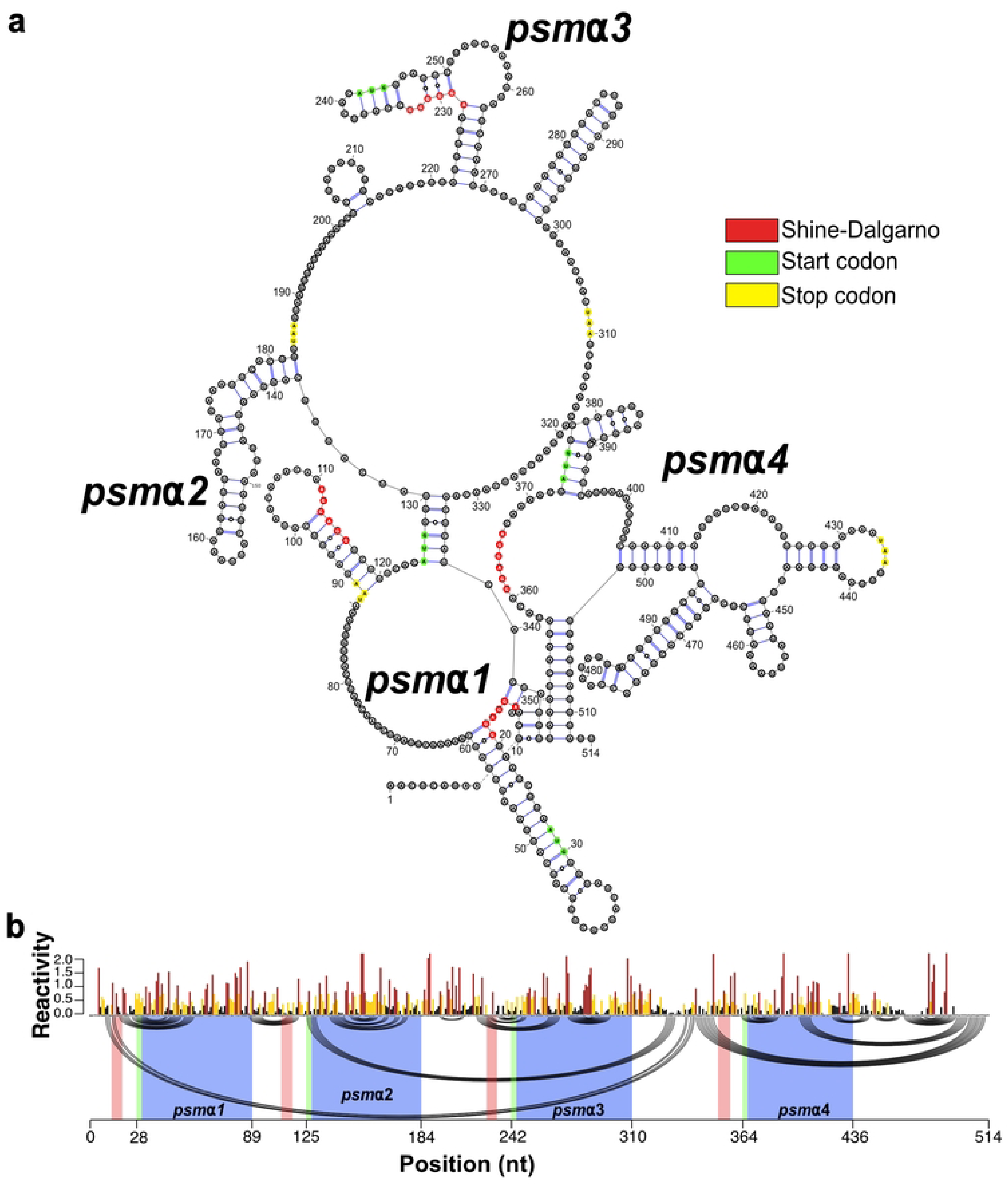
The *psm⍺* transcript is highly structured. **(a)** SHAPE-MaP-guided RNA secondary structure prediction of the *psm⍺* transcript in the parental USA300 strain. The Shine-Dalgarno (SD) sequence is marked in red, the start codon in green and stop codon in yellow. **(b)** SHAPE-Map reactivity values mapped onto the predicted base pairs to highlight structural differences. The Shine-Dalgarno (SD) sequence is marked in red, the start codon in green and the coding sequence in blue.

Thus, our SHAPE-MaP data also corroborate the prediction of a highly structured *psm⍺* transcript and suggest that *psm⍺4* may be preferentially translated due to its relatively accessible SD sequence.

### Disrupting RsaE increases ribosome occupancy on *psm⍺1*, *psm⍺2* and *psm⍺4* but not *psm⍺3*

Given the marked increase in *psm⍺* promoter activity in the *ΔrsaE* mutant and the known regulation of this promoter by AgrA from *agr* quorum sensing system [30,31], we hypothesised that RsaE deletion (directly or indirectly) enhances AgrA expression. To test this, we analysed both USA300 (Table S1) and published HG003 RNA sequencing data ([12] Table S2). Additionally, we analysed ribosome profiling datasets comparing ribosome occupancy in HG003 versus its *ΔrsaE* derivative ([12] Table S2).

Collectively, these data revealed that while *agrBDCA* transcript levels remained unchanged between parental vs. *ΔrsaE* strains in both USA300 (grown in TSB, Fig 4a) and HG003 *ΔrsaE* [12] backgrounds, ribosome occupancy on *agr* coding sequences was significantly higher in HG003 *ΔrsaE* vs. its parental strain.

**Fig 4.**
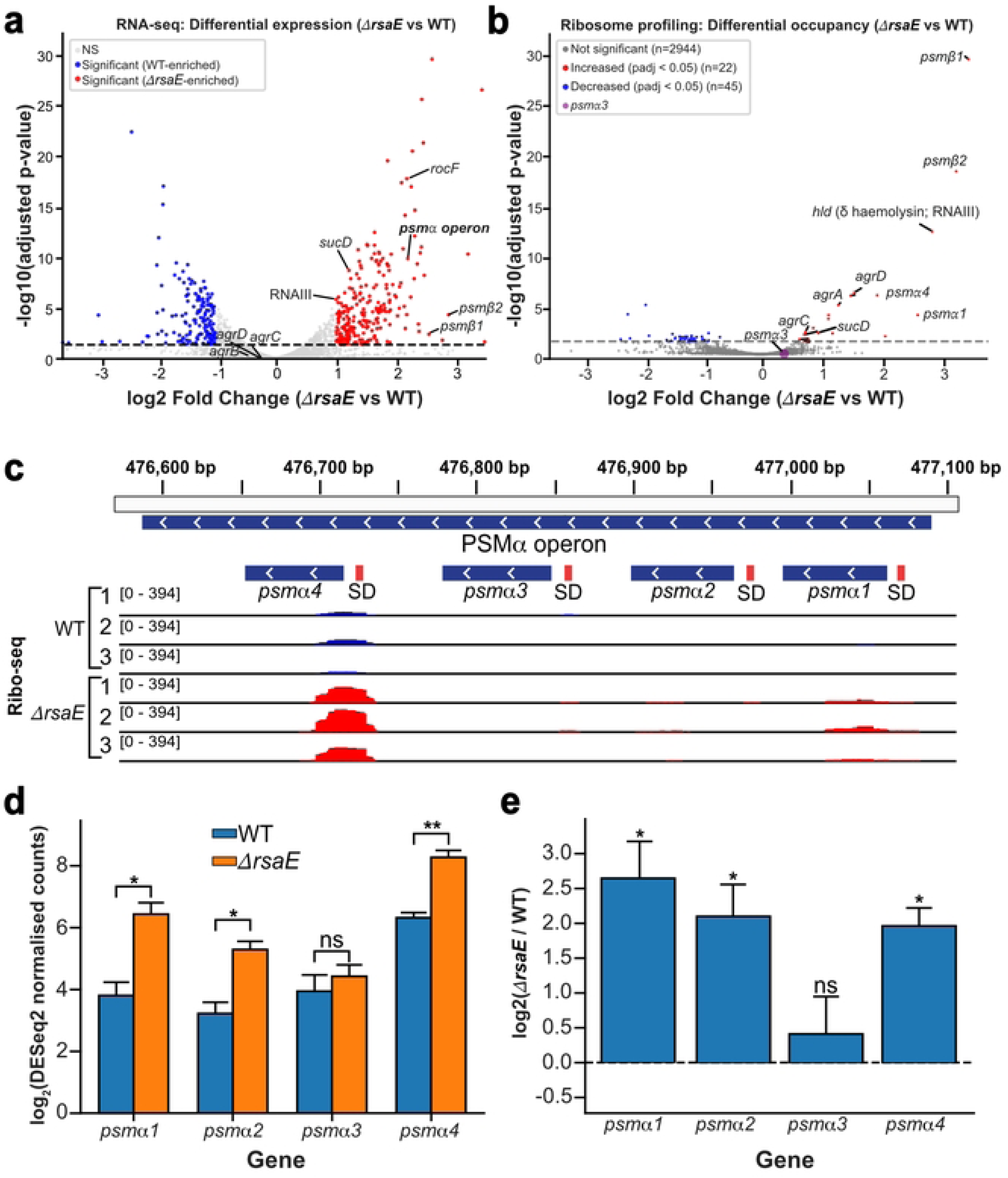
Deletion of RsaE activates the *agr* quorum sensing system. **(a)** Volcano plot showing transcriptome-wide changes in RNA steady-state levels between USA300 *ΔrsaE* and parental (WT) strain based on RNA-sequencing data. The x-axis represents log2-fold change in DESeq2 normalised read counts (*ΔrsaE* vs. WT), and the y-axis shows statistical significance as -log10(adjusted p-value). Gray dots indicate transcripts with no significant change. Red dots represent genes with significantly increased steady-state in the *ΔrsaE* mutant (padj < 0.05, n=22), while blue dots indicate genes with significantly decreased ribosome occupancy (padj < 0.05, n=45). The horizontal dashed line marks the significance threshold (padj = 0.05). Key genes are labelled, including *agr* quorum sensing components (*agrA, agrC, agrD*), PSM⍺ toxin operon, PSMβ toxins (*psmβ1*, *psmβ2*), delta-haemolysin (RNAIII), and *sucD*, a known RsaE target [13]. **(b)** Volcano plot showing transcriptome-wide changes in ribosome occupancy between HG003 *ΔrsaE* and parental (WT) strains. The x-axis represents log2-fold change in ribosome occupancy (*ΔrsaE* vs. WT), and the y-axis shows statistical significance as -log10(adjusted p-value). Gray dots indicate transcripts with no significant change in ribosome occupancy (n=2944). Red dots represent genes with significantly increased ribosome occupancy in the *ΔrsaE* mutant (padj < 0.05, n=22), while blue dots indicate genes with significantly decreased ribosome occupancy (padj < 0.05, n=45). The horizontal dashed line marks the significance threshold (padj = 0.05). Key genes are labelled, including *agr* quorum sensing components (*agrA, agrC, agrD*), PSM⍺ toxins (*psm⍺1*, *psm⍺3*, *psm⍺4*), PSMβ toxins (*psmβ1*, *psmβ2*), delta-haemolysin (hld/RNAIII), and again *sucD*. **(c)** Ribosome profiling coverage across the *psm⍺* operon locus. Ribosome-protected fragment (RPF) reads from three biological replicates are shown for wild-type (WT, blue) and *ΔrsaE* mutant (red) strains. The *psm⍺* operon is on the Crick strand and its structure is shown at the top with Shine-Dalgarno (SD) sequences marked in red. Results from three biological replicates for each strain are shown. **(d)** Quantification of ribosome occupancy on individual *psm⍺* genes. Bar graph shows log_2_-transformed DESeq2-normalized read counts for each *psm⍺* gene in WT (blue) and *ΔrsaE* (orange) strains. Three biological replicates were analysed. Error bars represent standard deviation. Statistical significance (adjusted p-values) was determined by DESeq. p < 0.05 = *, p < 0.01 = **, p < 0.0001 = ****. **(e)** Log2-fold change in ribosome occupancy between HG003 *ΔrsaE* and WT strains for individual *psm⍺* gene. Bars represent mean log_2_(*ΔrsaE*/WT) ratios with individual biological replicates shown as dots (n=3). Error bars represent standard deviation. Statistical significance was determined using a non-parametric Kolmogorov-Smirnov test. p < 0.05 = *, p < 0.01 = **, p < 0.0001 = ****.

Although our USA300 RNA-seq data were generated under different growth conditions (higher cell density,OD_595_=3.0), they align with the HG003 results [12] showing increased steady state levels of known RsaE mRNA targets in *ΔrsaE* (*sucC*, *sucD* and *rocF*; Fig 4a), but not *agr*. However, AgrA-regulated toxin genes (*psm⍺*, *psmβ* and *hld/*haemolysin 𝛿) showed significantly higher RNA steady state levels and ribosome occupancy in *ΔrsaE* (Fig 4b). These results imply substantial *agr* activation in *ΔrsaE*, predominantly through increased translation of its mRNAs, explaining the increased *psm⍺* promoter activity and elevated expression of toxin genes in *ΔrsaE*.

The ribosome occupancy on the *psm⍺* transcript (Fig 4c) corroborated numerous studies reporting differential expression of *psm⍺* toxins, with *psm⍺1 and psm⍺4* being consistently detected at higher levels [14,21,22]. This aligns with our SHAPE-MaP data, which predicted that *psm⍺4* would exhibit a higher ribosome occupancy, followed by *psm⍺1.* In contrast, *psm⍺2* and *psm⍺3* showed low levels of ribosome occupancy, supporting the hypothesis that sequestration of their AUG and SD sequences in stem structures impairs translation. Deleting RsaE resulted in a significant increase in ribosome occupancy on *psm⍺1*, *psm⍺2* and *psm⍺4* (Fig 4b-d), consistent with elevated *psm⍺* transcript levels. However, ribosome occupancy on *psm⍺3* remained unchanged (Fig 4d-e).

This result suggests that the increase in *psm⍺* operon expression upon *rsaE* deletion does not concomitantly increase ribosome occupancy on *psm⍺3*, suggesting that intrinsic SD-sequestering secondary structures play a dominant role in regulating *psm⍺3* translation. We therefore propose that RsaE base-pairing with the *psm⍺3* SD sequence may fine-tune its expression, rather than acting as a primary regulator.

### The 5’ UCCCC seed sequence of RsaE is necessary and sufficient for *psm⍺* regulation

To decipher how RsaE regulates the stability of the *psm⍺* transcript, we generated RsaE mutants with disrupted base-pairing seed sequences. RsaE contains two duplicated conserved C-rich regions (CRRs), one at the 5′ end and one near the 3′ end, either of which could potentially mediate pairing with the *psm⍺* SD sequence (Fig 5a) [12]. To evaluate the functional contribution of each motif *in vivo*, a series of complementation strains were generated in the USA300 *ΔrsaE* background. These strains expressed either wild-type RsaE, a 5′-motif mutant (mut-1), a 3′-motif mutant (mut-2), or a double mutant lacking both motifs (both; Fig 5a). In all mutants the C’s were substituted with G’s, preventing the motif from base-pairing with G-rich targets, such as SD sequences.

**Fig 5.**
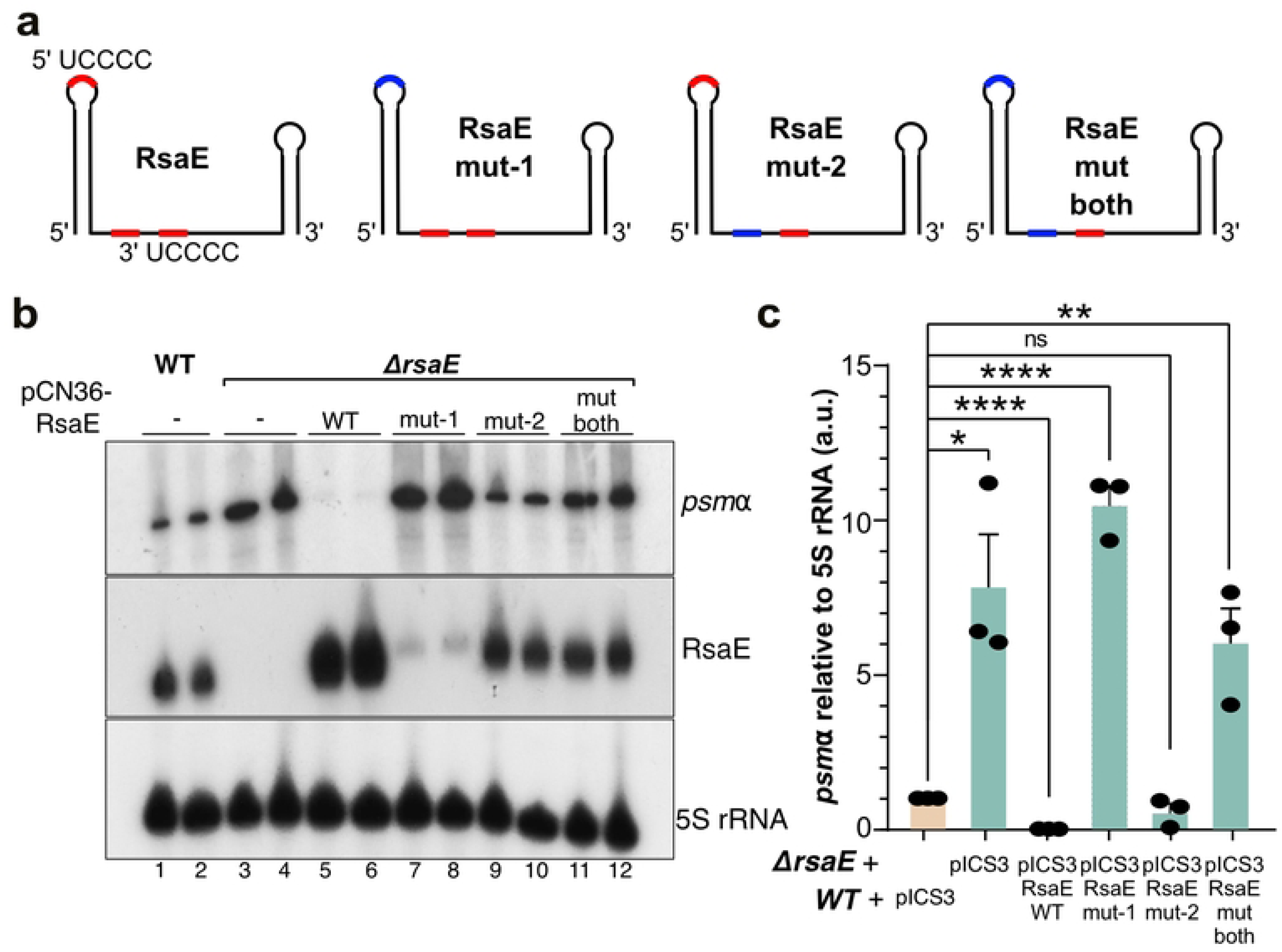
The 5′ seed sequence of RsaE regulates *psm⍺* expression. **(a)** Schematic representation of RsaE with its 5’ and 3’ seed sequences. The mutated seed sequence is shown in red. **(b)** Northern blot analysis of *ΔrsaE* strains complemented with wild-type or seed sequence mutant variants of RsaE (expressed from the pICS3 plasmid). The top panel was probed for *psm⍺*, the middle panel for RsaE, and the bottom panel shows 5S rRNA as a loading control. **(c)** RT-qPCR analysis showing fold change of *psm⍺* expression normalized to 5S rRNA across the same strains. Data represent the averages and standard error of the mean (SEM) from three biological replicates. Statistical significance was determined using an unpaired t-test (p < 0.05 = *, p < 0.01 = **, p < 0.0001 = ****).

Northern blotting confirmed expression of RsaE and allowed comparison of *psm⍺* transcript abundance across strains (Fig 5b). RT-qPCR was used to support these observations (Fig 5c). Northern blot experiments were performed with two biological replicate RNA samples for each strain. Consistent with our previous analyses, deletion of RsaE increased *psm⍺* steady state levels (Fig 5b, lanes 3-4 vs. 1-2; Fig 5c). Complementing the *ΔrsaE* strain with wild-type *rsaE* expressed from the pICS3 plasmid resulted in RsaE over-expression and the *psm⍺* transcript was almost undetectable (Fig 5b; lanes 5-6 vs. 1-2; Fig 5c). Hence, increasing RsaE levels results in increased decay of *psm⍺* RNA and/or reduced *psm⍺* transcription. Unfortunately, the mut-1 RsaE construct (first CRR mutated) was unstable and therefore no conclusions can be drawn about this mutant (Fig 5b; lanes 7-8; Fig 5c). The mut-2 (second CRR mutated) and mut-both (both CRRs mutated) were expressed at levels comparable to wild-type RsaE (Fig 5b, lanes 9-12 vs. 1-2). In the strain expressing the mut-2 mutant, *psm⍺* transcript levels were comparable to those in the parental strain (Fig 5b, lanes 9-10 vs. 1-2; Fig 5c). Thus, this mutation does not significantly impact RsaE-mediated decay of *psm⍺*. In contrast, mutating both seed sequences in RsaE generated similar results as the non-complemented *ΔrsaE* strain (Fig 5b; lanes 11-12 vs. 1-2; Fig 5c). While these results do not rule out a role for the 3’-UCCCC RsaE sequence in regulating decay of the *psm⍺* transcript, we conclude that the 5’-UCCCC motif is necessary and sufficient for interacting with *psm⍺*.

### RNase III is required for *in vitro* biofilm formation in USA300

To assess the biological impact of the increased *psm⍺* transcript levels in the *ΔrsaE* and *Δrny* mutants, we investigated their effects on biofilm formations, given the importance of PSM⍺ toxins in regulating biofilm structuring, stability and dissemination [32]. Before starting this experiment, we verified that none of the gene deletions affected cell growth, as alterations in growth rates could influence biofilm production or stability. Growth curve analyses revealed no statistically significant differences in growth rates between the mutants and the parental USA300 strain (S4 Fig).

For analysing biofilm formation, cells were grown in TSB medium in 96 well plates and biofilm production was monitored for 24, 48 and 72 hours. After this time, the planktonic cells were gently removed, biofilms were fixed with formaldehyde and stained with crystal violet (CV) that binds to negatively charged molecules, such as nucleic acids and proteins. CV retention measured by absorbance at 595 nm after acetone treatment, served as a proxy for the biofilm biomass. We also included *Δrnc* and *Δpsm⍺* strains in this analysis.

Much to our surprise, the *Δrnc* strain already showed a markedly reduced biofilm biomass compared to all other strains (Fig 6a-b). This persisted across biological replicates and 48- and 72-hour time points, indicating that RNase III contributes to both early biofilm establishment, such as surface attachment, and to its continued maintenance. To our knowledge, this is the first evidence linking RNase III to biofilm formation in methicillin-resistant *S. aureus*.

**Fig 6.**
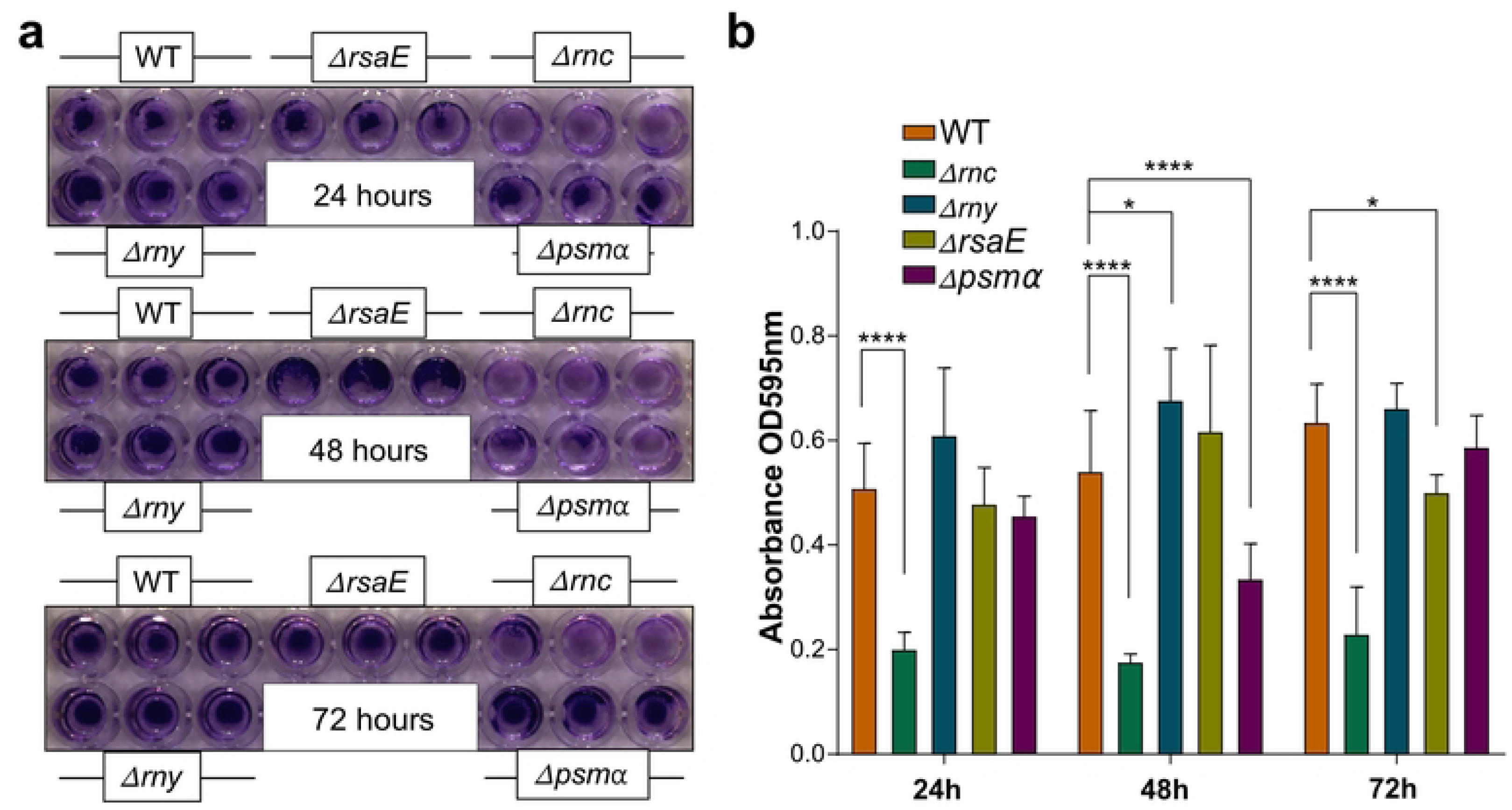
RNase III is required for biofilm formation. **(a)** Images of biofilm formation by each mutant strain stained with Crystal Violet after 24, 48, and 72 hours of static incubation. **(b)** Quantification of biofilm biomass by measuring CV absorbance at OD_595_ after diluting the solubilised Crystal Violet in acetic acid (1:6 dilution). Strains are indicated along the top of the bar plot. Statistical significance was assessed using two-way ANOVA followed by post hoc Dunnett test (p < 0.05 = *, p < 0.001 = ***, p < 0.0001 = ****)

The biofilms formed by the *Δpsm⍺* mutant showed less CV staining at 48h, but this was largely restored after 72 hours. The *Δrny* mutant showed no major differences in biofilm formation in the CV assay compared to the USA300 parental strain across most time points. However, a modest but statistically significant increase in staining was observed at 48 hours (Fig 6b), suggesting increased biomass. Deletion of *rsaE* had no noticeable impact on biofilm biomass during the first 48 hours, but a significant decrease in biomass was observed at the 72-hour time point (Fig 6b).

We conclude that the increased *psm⍺*transcript levels observed in *ΔrsaE* and *Δrny* mutants only modestly alter USA300 biofilm CV staining under the tested *in vitro* conditions.

### RsaE regulates USA300 biofilm thickness and eDNA layer structuring *in vitro*

To complement the semi-quantitative data obtained from CV assay, confocal laser scanning microscopy (CLSM) was employed to directly visualise biofilm architecture at the microscale level within microtiter wells. SYPRO Ruby was used to stain proteins. In *Staphylococcus epidermidis*, RsaE promotes biofilm structuring by increasing Poly-N-acetylglucosamine (PNAG) levels, likely by inducing the *icaADBC* operon through repression of the repressor *icaR*. RsaE has also been shown to base-pair with *icaR* in *S. aureus* and regulate its steady state levels [12]. Moreover, *S. epidermis* RsaE regulates extracellular DNA (eDNA) release in biofilms [33]. To assess whether *S. aureus* RsaE similarly influences biofilm matrix composition, we used wheat germ agglutinin (WGA) conjugated with Alexa Fluor 488 to detect PNAG. To quantify eDNA release, TOTO-3 iodide was used to detect the B-form DNA and the Z22 rabbit monoclonal IgG kappa antibody to detect Z-DNA. Monitoring Z-DNA accumulation in biofilms provides information about the maturation state, as Z-DNA is the predominant structural form of eDNA in mature biofilms [34]. Finally, Hoechst dye staining of DNA was used as a proxy for biomass. Since the cells were not permeabilised, intense Hoechst staining indicated likely dead cells.

Combined, these staining approaches allowed for the comparative assessment of biofilm matrix production and biofilm morphology across different genetic backgrounds and developmental stages.

Biofilms were cultivated up to 24, 48 and 72 hours prior to analysis (Fig 7). For the eDNA analysis only biofilms grown up to 72 hours were assessed (Fig 8), as this time point was expected to coincide with nutrient-depletion-induced stress, a condition associated with increased Z-DNA formation in eDNA for several bacterial species [34]. We analysed images from two biological replicates containing 2-4 technical replicates each. To quantify cellular signal intensities and cell counts, we developed a custom Python image analysis pipeline utilising the Cellpose deep learning cell segmentation Python API ([35] see Data availability). The pipeline performs automated segmentation of individual bacterial cells across multi-channel confocal Z-stacks and quantifies fluorescence intensities on a per-cell basis, as outlined in the Methods section. Representative images of the microscopy analyses and the result of the quantification analyses are shown in S5 Fig, Fig 7 and Fig 8. The results of the statistical analysis of data are provided in S6 Fig.

**Fig 7.**
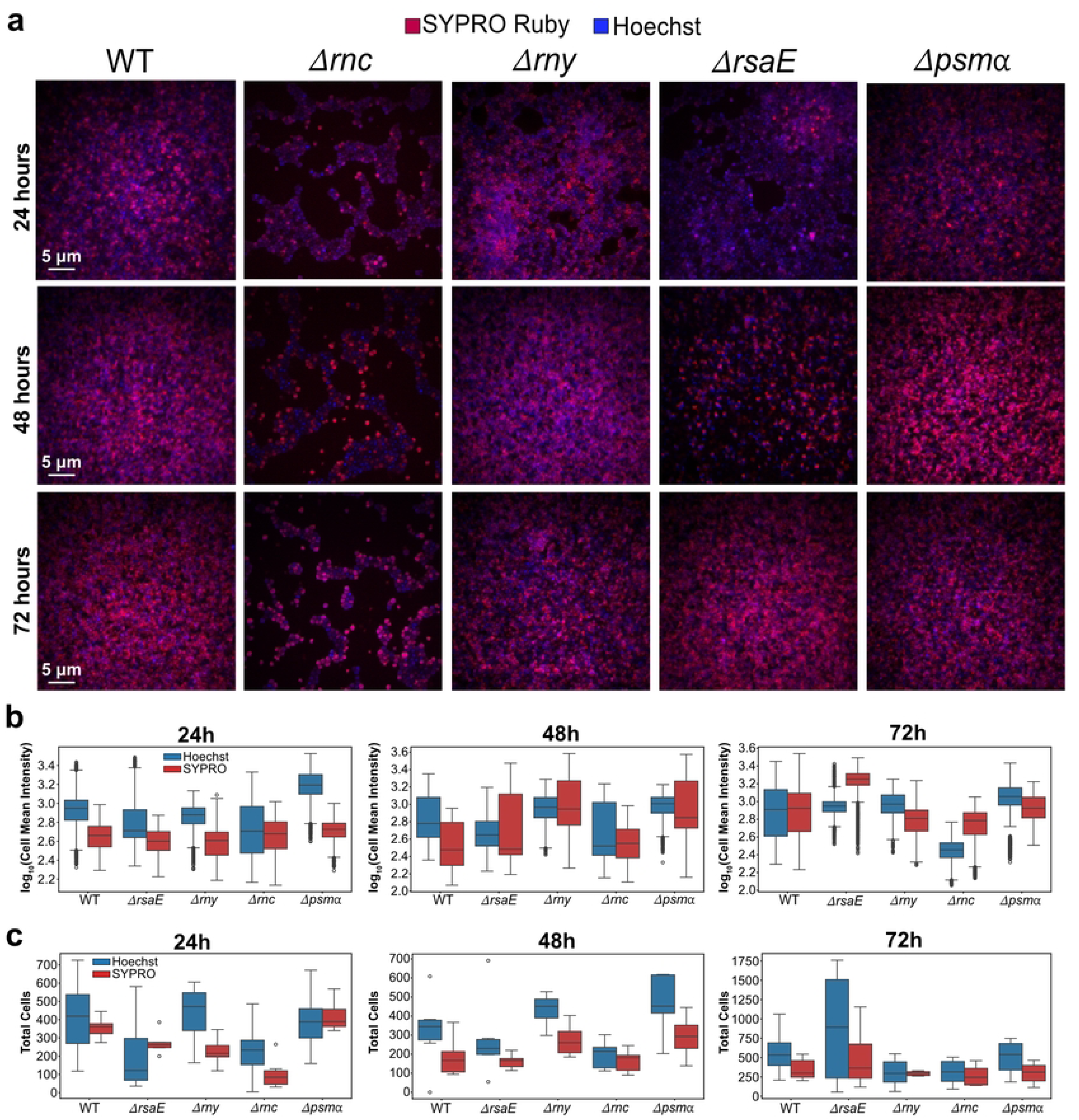
Analysis of biofilm mass differences between the parental USA300 strains and deletion mutants. **(a)** Biofilms were imaged after static incubation using confocal laser scanning microscopy. The following fluorescent stains were used: Hoechst (blue) for nucleic acids (excitation: 405 nm, emission: 447 nm) and SYPRO Ruby (red) for extracellular proteins (excitation: 561 nm, emission: 561 nm). Shown are representative image from one technical replicate. In total 2 biological replicates and 2-4 technical replicates for each biological replicate were analysed. Scale bar: 5 µm. **(b)** Quantification of protein and DNA content at the 24, 48 and 72-hour time point using our Cellpose Python pipeline (see Data availability). Box plots display log_10_-transformed cell mean intensity values for each channel, based on two biological replicates with 2-4 technical replicates for each strain. **(c)** Quantification of the number of cells stained with Hoechst and SYPRO dyes at the 24, 48 and 72-hour time point using our Cellpose Python pipeline. Error bars indicate standard deviations obtained from two biological replicate samples with each 2-4 technical replicates.

**Fig 8.**
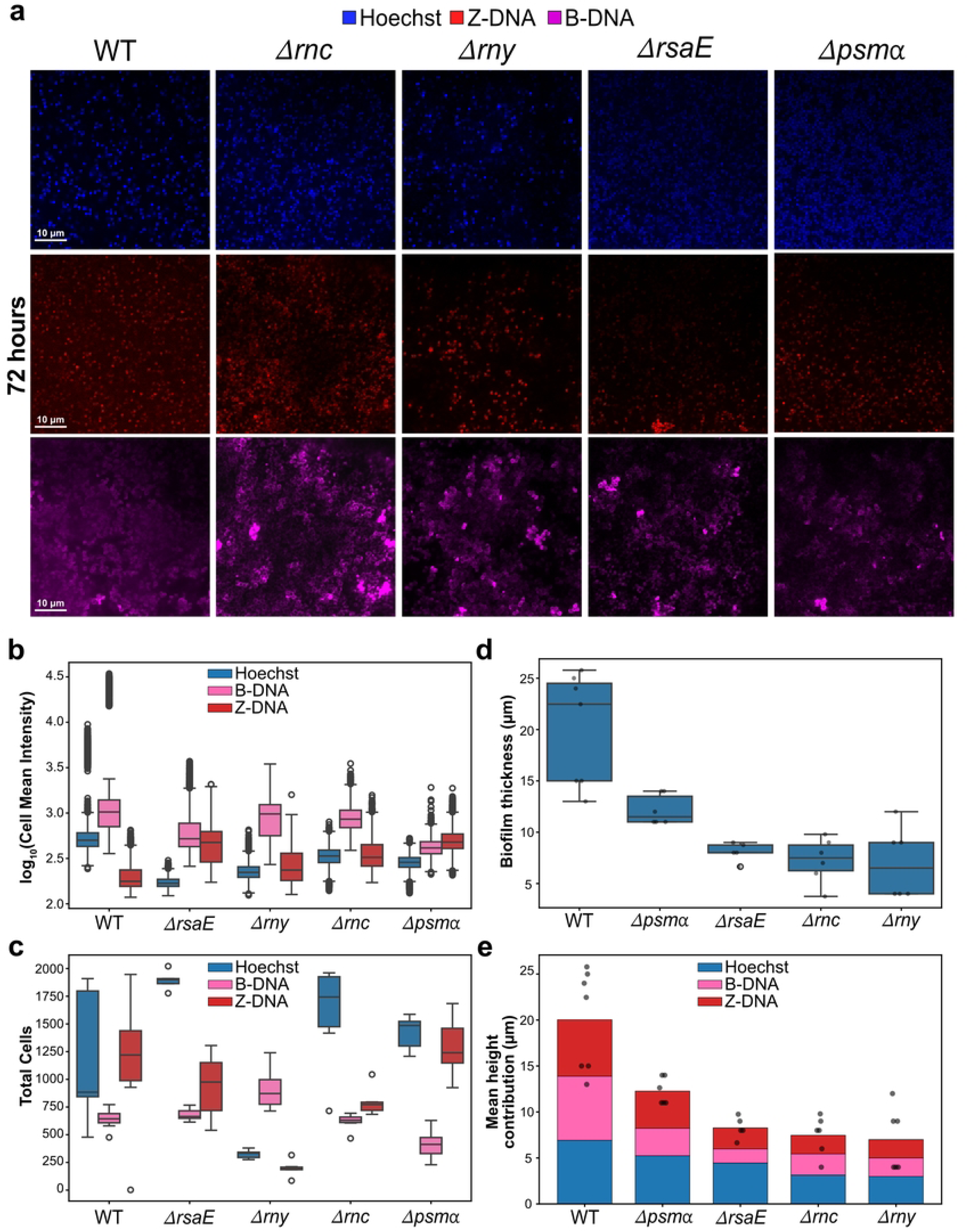
RsaE regulates biofilm thickness and eDNA layer formation. **(a)** Biofilms were imaged after static incubation up to 72 hours using confocal laser scanning microscopy. The following fluorescent stains were used: Hoechst (blue) for nucleic acids (excitation: 405 nm, emission: 447 nm), TOTO-3 (magenta) for B-DNA (excitation: 561 nm, emission: 630 nm), and anti-ZDNA Z-22 Rabbit IgG antibody followed by secondary Donkey Anti-Rabbit antibody conjugated with Alexa Fluor 594 (red) for eDNA:Z-DNA (excitation: 640 nm, emission: 660 nm). Scale bar: 10 µm. **(b)** Quantification of (e)DNA content at the 72-hour time point using our Cellpose Python pipeline (see Data availability). Box plots display log_10_-transformed cell mean intensity values for the Hoechst and l, based on two biological replicates each with 2-4 technical replicates for each strain. Error bars indicate standard deviations obtained from two biological replicate samples with each 2-4 technical replicates. **(c)** Quantification the number of cells with (e)DNA signals at the 72-hour time point using our Cellpose Python pipeline. Error bars indicate standard deviations obtained from two biological replicate samples with each 2-4 technical replicates. **(d)** Biofilm thickness of *S. aureus* USA300 and indicated deletion strains at 72h of growth. Biofilm thickness was measured per field of view as the Z-distance between the shallowest and deepest segmented cell centroid, taking the union of the Hoechst (blue) and TOTO-3 (magenta) channels. Each dot represents one independent field of view (technical replicate). Boxes show the median and interquartile range; whiskers extend to 1.5× IQR. Statistical comparisons against WT were performed using the two-sided Mann–Whitney U test (WT n = 7 fields of view; *Δpsm* n = 6; *Δrnc* n = 6; *Δrny* n = 6; *ΔrsaE* n = 5). All deletion strains showed significantly reduced biofilm thickness compared to WT: *Δpsm* median 11.5 µm (p = 0.008); *Δrnc* median 7.5 µm (p = 0.003); *Δrny* median 6.5 µm (p = 0.003); *ΔrsaE* median 8.0 µm vs WT median 22.5 µm (p = 0.006). All comparisons remain significant after Bonferroni correction for four tests (adjusted threshold p < 0.0125). **(e)** Per-channel contributions to biofilm height across genotypes at 72h of growth. Stacked bars show the mean contribution of each fluorescence channel to total biofilm height, averaged across all fields of view for each genotype. For each field of view, the Z-span of each channel (distance between the shallowest and deepest segmented cell centroid) was computed; contributions were calculated as the proportional Z-span of each channel relative to the sum of all channel spans, scaled by total biofilm height, such that bar segments sum to the total height. Colours indicate fluorescence channel: blue, Hoechst; magenta, TOTO-3 (B-DNA); red, Z-DNA. Black dots show the total measured biofilm height for each individual field of view. Data are from the 72h Z-DNA dataset (2 biological replicates, 2–3 fields of view each).

To ensure that analysis was restricted to in-focus biofilm-containing regions, the pipeline computed a signal-to-noise ratio (SNR) metric across Z-slices for each channel and identified the primary focus peak using Gaussian fitting to the smoothed SNR trace (S7b Fig). Following focus selection, individual cells were segmented using the pre-trained (cyto2 model) in 2D+stitch mode, in which 2D segmentation masks from consecutive Z-slices are stitched across Z by IoU-based label propagation. Post-segmentation refinement was performed to enforce biologically plausible cell morphologies through roundness constraints and volume-based filtering. Statistical significance between each mutant and the WT was evaluated using Mann-Whitney U combined with Benjamini-Hochberg correction. Because the high sample size could bias the calculation of *p*-values [36], the Cohen’s *d* test [37] was also used to also assess the effect size.

The WGA staining of USA300 biofilms generated unexpected results and prompted us to test the specificity of this dye on our biofilms. For this purpose, included a *ΔicaADBC* transposon mutant as a negative control, as it does not produce PNAG. A *ΔicaR* transposon mutant that produces high levels of PNAG was used as positive control (S5a Fig). Incubating the USA300 biofilms with WGA indicated mostly non-specific binding of the dye: WGA staining could readily be detected in all tested stains by microscopy, including our negative control strain (S5a Fig; *ΔicaADBC*). These data imply cross-reactivity of WGA with peptidoglycans in the cell wall. During the troubleshooting of these results, we quantified the *icaR* and *icaA* steady-state transcript levels in our strains (S5c-d Fig). Illumina and Nanopore RNA sequencing analyses of the parental strain grown in the TSB medium used for growing biofilms (TSB) showed that while *icaR* was abundantly expressed, *icaABC* could only be detected at low levels (S5b Fig). Moreover, qRT-PCR and ribosome profiling data showed that knocking out RsaE had no significant impact on *icaR* or *icaADBC* expression and ribosome occupancy (S5c-e Fig). Thus, when USA300 is grown in the nutrient rich TSB medium we used for growing biofilms, the expression levels of RsaE are too low to have a meaningful impact on *icaR* and *icaADBC*. Only when RsaE was overexpressed from the pICS3 plasmid could a significant decrease in *icaR* expression (S5c Fig) and a concomitant increase in *icaA* levels be observed (S5d Fig).

Given these results, we concluded that WGA staining is not suitable to quantify PNAG production in USA300 biofilms. Moreover, PNAG synthesis was very low under the biofilm growth conditions used, suggesting it is highly unlikely to have a meaningful impact on biofilm formation in this strain under the tested growth conditions, similarly to what was observed in other MRSA strains [38]. Hence, while we did perform the WGA imaging on all strains (see Data Availability), we did not consider these results further. Nevertheless, this investigation did reveal that, under the growth conditions used, RsaE does not accumulate at sufficient levels to significantly alter PNAG synthesis.

We next used our Cellpose pipeline to count the total number of cells in each biofilm as well as the mean signal intensity of each dye in each individual cell. At the 24-hour time point, the wild-type (WT) strain exhibited comparably distributed fluorescent signals across both protein (red) and bacterial DNA (blue) channels (Fig 7a-b; S6a Fig). This was sustained at later time points. Consistent with our CV staining data (Fig 6), the *Δrnc* mutant biofilms displayed lower cell counts as well as lower signal intensities with both Hoechst and SYPRO staining. Cohen’s d effect sizes indicated that differences in cell counts and signal intensities in both channels were substantial after 24 hours (Fig 7a-c, S6a-c Fig).

The *Δrny* strain exhibited cell counts and fluorescence intensities comparable to, or slightly higher than, WT (Fig 7a-c; S6a-c Fig). However, as with the *Δrnc* mutant, the total cell counts for the Hoechst channel in the *Δrny* mutant were significantly lower compared to WT after 72 hours of growth (Fig 7a-c; S6a-c Fig). This implies that RNase Y is required for maintenance of biofilm mass at later stages of biofilm growth. The *ΔrsaE* biofilms looked similar to WT at the 24h and 48h time points; however, we note lower counts of cells stained with Hoechst at these time points. At 72h, the cell counts were highly variable for the *ΔrsaE* strain, making it difficult to draw conclusions. The *Δpsm⍺* mutant showed higher cell counts in biofilms in both Hoechst and SYPRO channels at 48-hour time point. This is consistent with previous work showing increased biofilm volume in USA300 LAC *Δpsm⍺* [39]. However, at the 72-hour time point, biofilm cell counts were comparable to the parental strain.

Biofilm mass and thickness are not only determined by cell number and protein content, as shown by our Hoechst and SYPRO staining results, but also by extracellular DNA (eDNA), which can make a significant contribution to biofilm architecture and thickness. We therefore next investigated the abundance and conformational state of eDNA in the biofilms after 72h of growth (Fig 8; S6d Fig). The bulk of the eDNA signals was closely associated with the bacterial cell wall rather than appearing as a diffuse matrix (Fig 8a). Therefore, we focussed our analyses on cell-surface associated eDNA. Importantly, the B- and Z-DNA signals were distinct from total DNA (Hoechst), localising as structures above the Hoechst-stained bacterial cells. This is consistent with matrix-associated extracellular DNA staining (S7a-c Fig).

Our analyses revealed notable differences in biofilm structure and thickness in the mutants compared to WT. Consistent with the results shown in Fig 7, we note that the *Δrny* strain showed a significantly lower number of cells with Hoechst, but also extracellular Z-DNA staining (Fig 8a-c; S6d Fig). B-DNA signals, however, were significantly higher in this strain, suggesting a defect in biofilm maturation. Contrary to the results shown in Fig 7, in this experiment the number of cells stained with Hoechst in the *Δrnc* biofilms was comparable to WT; however, we did observe a lower number of cells displaying B-DNA and Z-DNA signals. This discrepancy in cell counts between WT and *Δrnc* may be attributable to the many additional incubation and washing steps required for Z-DNA detection, which may have dislodged some of the biofilm. Indeed, we noted higher variability in cell counts across replicate biofilms, in particular with the parental (WT) strain (Fig 8c).

Nevertheless, the most illuminating findings came not from cell counts but from measurements of biofilm thickness and eDNA architecture. By far the most striking differences between the mutants and the parental strain were in the thickness of the corresponding biofilms and the contribution of the eDNA layer to biofilm structure. This is evident from Figs 8d and e, which show markedly reduced biofilm thickness and a diminished contribution of eDNA to overall biofilm depth across the mutant strains. The 3D Cellpose renderings of the biofilms revealed that WT biofilms showed a dense, uniform signal in both eDNA channels, reflected in the tall, even surface in the 3D plots (S8b Fig). Strikingly, although in many cases the cell counts for each eDNA channel were not substantially different from WT, the thickness of the eDNA layer was noticeably reduced in all deletion mutants. Notably, in the *Δrny* and *ΔrsaE* biofilms, eDNA appeared to be more evenly distributed throughout the biofilm rather than concentrated at its surface. This suggests that RsaE and RNase Y influence the spatial organisation of eDNA within the biofilm matrix.

### RsaE regulates *in vitro* biofilm viability

In addition to regulating toxin expression, RsaE represses genes involved in the TCA cycle. Therefore, we examined whether de-repression of the TCA cycle in the *ΔrsaE* mutant affects the metabolic fitness and viability of cells in biofilms. We also included the other deletion strains described above in the assay. To determine the number of viable cells within biofilms formed by different mutants, initial CFU assays were performed at 24, 48, and 72 hours. The wild-type strain showed intermediate CFU levels, with a modest decrease over time. This reflects a balance between biofilm formation and cell survival, serving as a baseline for comparison with the mutant phenotypes.

The general trend observed across most mutant strains was a gradual decline in viable cell counts over time. Fig 9a illustrates the viability trends across all mutant strains. The *Δrnc* strain exhibited intermediate CFU counts with even more live cells at 48 hours than the WT. This is noteworthy given the strain’s reduced biofilm-forming ability, indicating that although biofilm architecture is compromised, the viability of the remaining bacterial population remains relatively high. The *Δrny* mutant showed viability results comparable to the parental strain. The *Δpsm⍺* mutant, however, had the highest CFU count after 24 hours of biofilm growth, but showed a sharp decline after 48 hours across biological replicates, suggesting that deletion of *psm⍺* creates conditions within the biofilm that does not support sustained bacterial proliferation and survival.

**Fig 9.**
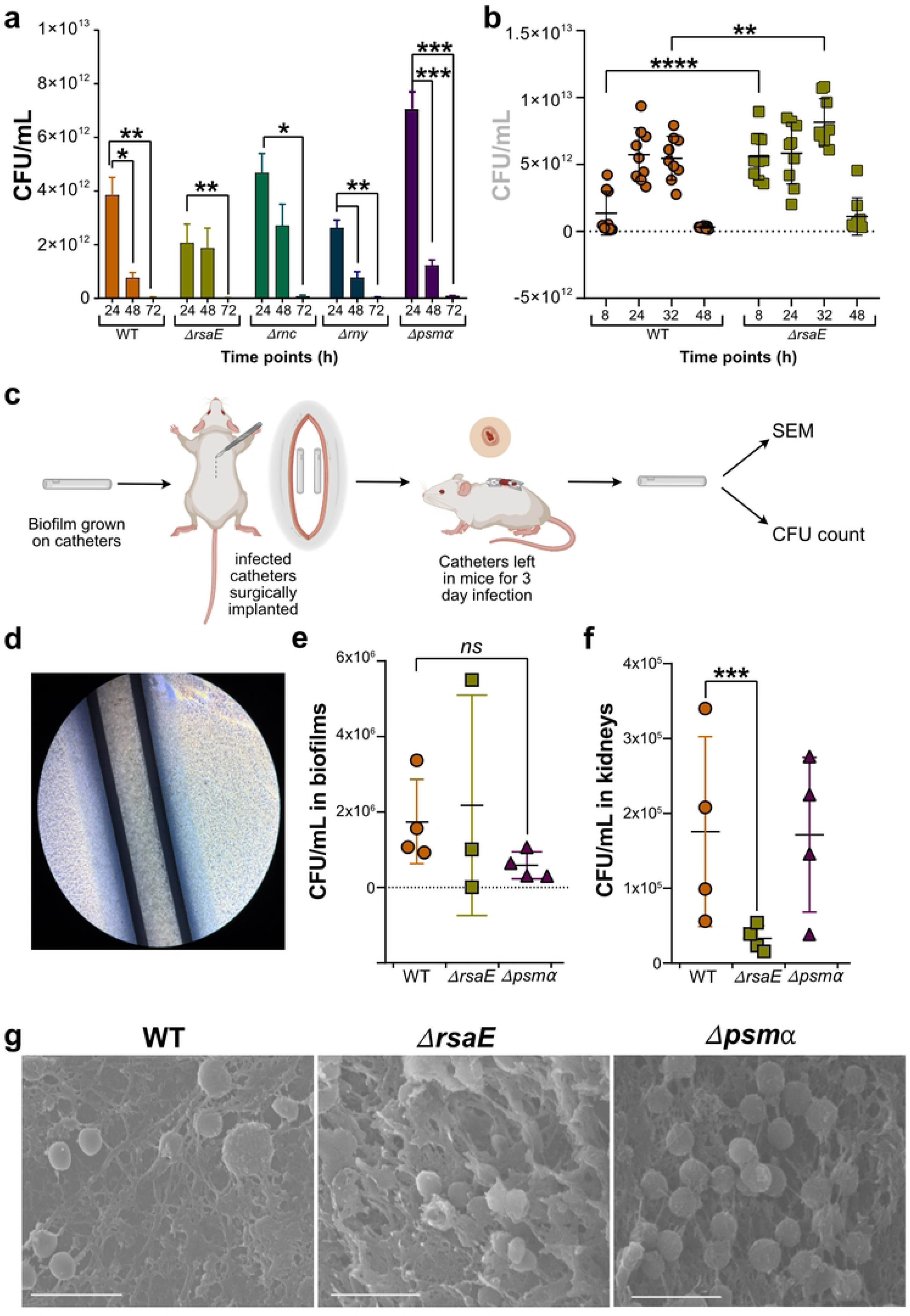
RsaE regulates biofilm viability and dissemination. **(a)** Bar plot showing colony-forming units per millilitre (CFU/mL) for each strain at 24, 48, and 72 hours of biofilm growth. Data represent the mean of three biological replicates, each with three technical replicates. Statistical significance was determined using two-way ANOVA with Sidak’s multiple comparison (p < 0.05 = *, p < 0.001 = ***, p < 0.00001 = ****). **(b)** Individual CFU/mL values for wild-type and *ΔrsaE* strains plotted at 8, 24, 32, and 48 hours. Statistical significance was assessed using two-way ANOVA with with Sidak’s multiple comparison (p < 0.05 = *, p < 0.001 = ***, p < 0.00001 = ****) **(c)** Schematic illustrating the procedure for placing a catheter under the skin of mice followed by CFU analysis and imaging. Illustration was made in BioRender. **(d)** Catheter segments with 24-hour *in vitro* biofilms visualised under a light microscope prior to implantation. **(e)** Individual CFU/mL values recovered from wild-type and *ΔrsaE* catheters after 3-day infection. **(f)** Individual CFU/mL values recovered from wild-type and *ΔrsaE* kidneys after 3-day infection. **(g)** SEM images of explanted catheters colonised by wild-type, *ΔrsaE*, and *Δpsm⍺* strains at days 3 post-infection. All scale bars represent 5 µm.

The *ΔrsaE* mutant consistently maintained higher levels of viable cells until the 48-hour time point, although the data for the 24-hour time point were more variable and not significantly different from WT. These data suggest a potential role for RsaE in modulating bacterial survival or burden within the biofilm environment under stress conditions. To investigate this in more detail, additional CFU assays were conducted with *ΔrsaE* at earlier and intermediate time points: 8, 12-, 24-, 32-, and 42-hours using WT as a baseline control. This expanded time course provided a more detailed profile of bacterial viability in the absence of RsaE. Except for the 24-hour time point, the *ΔrsaE* mutant maintained slightly higher CFU levels than the wild-type especially being statistically significant at 8- and 32-hour time points. This implies that the bacteria in the *ΔrsaE* biofilms benefit from altered metabolic states that enhance survival during early biofilm growth phases. These results provide an important layer of context to the structural data from microscopy and crystal violet assays. They demonstrate that biofilm robustness, structure and bacterial viability do not always correlate directly.

We conclude that RsaE acts as a metabolic restraint that limits MRSA biofilm cell viability during establishment and mid-phase development, with regulatory effects that are ultimately overcome by late-stage biofilm stresses.

### RsaE enhances *S. aureus* dissemination from *in vivo* biofilms during early infection stages

Since the absence of RsaE increased the proportion of viable cells within biofilms in vitro without markedly altering biomass and structure, we sought to determine whether this effect extended to an *in vivo* setting. While *in vitro* biofilm studies in rich media (TSB) are informative for dissecting molecular mechanisms, they may not recapitulate the complex host environment where nutrient limitation, immune pressure, and tissue interactions fundamentally alter bacterial physiology. Therefore, validation in an *in vivo* infection model is important to understand the true functional role of RsaE in biofilm-associated disease. We reasoned that a higher number of surviving cells within the matrix could enhance progression of infection in the host. To test this, we employed an *in vivo* murine catheter associated biofilm model using wild-type, *ΔrsaE*, and *Δpsm⍺* mutant strains. The latter strain was included as previous work had shown that deleting this operon reduced biofilm dissemination in a similar mouse catheter model after seven days of infection [39]. Mice were subcutaneously implanted with sections of catheter tubing that had been inoculated with each strain, thus allowing biofilms to form on the catheters *in vivo* (Fig 9c). Based on preliminary experiments, catheters and kidneys were harvested at 3 days post-infection, a time point that captured early biofilm development and dissemination while avoiding confounding effects from advanced systemic/surgical wound infection. Prior to implantation, catheters were examined under a light microscope to ensure the presence of pre-formed biofilms, confirming consistency across experimental groups (Fig 9d). Upon euthanasia, catheters were explanted for CFU assays (Fig 9d-f) and scanning electron microscopy (SEM; Fig 9g). Kidneys were taken for assessing dispersal of the bacteria from the catheter biofilms (Fig 9f).

The *in vivo* data revealed a phenotype distinct from our *in vitro* observations. While *ΔrsaE* showed elevated cell viability in TSB biofilms (Fig 9b), the bacterial load in kidneys was significantly lower compared to both the parental strain and the *Δpsm⍺* mutant (Fig 9f), indicating reduced dissemination from catheter biofilms during early infection. Moreover, SEM analysis revealed that *ΔrsaE* biofilms were structurally distinct *in vivo* compared to the *in vitro* phenotype (Fig 9g), with sparser bacterial colonisation and gelatinous matrix production that was not evident in laboratory culture conditions. Wild-type biofilms on the catheters were characterised by spherical cocci beginning to cluster. In contrast, *ΔrsaE* biofilms appeared structurally distinct after three days of infection. This included sparser bacterial colonisation and gelatinous matrix production. The *Δpsm⍺* strain exhibited yet another phenotype. The matrix production was more spider web like and bacterial clustering was present.

Together, both the *in vitro* and *in vivo* findings underscore the importance of RsaE in shaping biofilm development, host interaction, and infection outcomes.

## Discussion

### RsaE regulates *psm⍺* toxin transcript levels at the transcriptional and post-transcriptional level

Using the CLASH approach with RNase III as bait, we previously uncovered large RNA-RNA interactomes in two clinically relevant MRSA strains [14,40]. These experiments identified known RsaE-RNA target interactions as well as several novel putative RsaE regulatory targets, including transcripts encoding the PSM⍺ and PSMβ toxins, short peptides that play key roles in biofilm structuring, dissemination, and immune evasion. We confirmed the RsaE::*psm⍺* transcript base-pairing *in vitro* [14], however, the biological significance of this interaction remained unclear.

To investigate the functional role of RsaE in toxin regulation, we examined gene expression in a *ΔrsaE* deletion strain and compared it to the parental strain. Deletion of RsaE resulted in increased steady-state levels of *psm⍺* transcripts as well as upregulation of *psmβ* and *hld* (encoding δ-toxin) mRNAs, demonstrating that RsaE has a broader impact on virulence gene expression than previously anticipated. RNA half-life measurements revealed that *psm⍺* transcripts are remarkably stable, with a mean half-life of approximately 15 minutes. SHAPE-MaP chemical probing of the *psm⍺* transcript revealed extensive secondary structure, including stem-loop elements that sequester the Shine-Dalgarno sequences of *psm⍺2* and *psm⍺3*. This provides a structural basis for both the exceptional transcript stability and the differential translation of individual PSM*⍺* peptides. Deleting RsaE further increased the *psm⍺* transcript half-life and elevated steady-state levels, suggesting a post-transcriptional role for RsaE in regulating the stability of this transcript. However, our data suggest that the increase in *psm⍺* steady-state levels in *ΔrsaE* is predominantly attributable to transcriptional activation of this operon via the *agr* quorum sensing system (see below). Our ribosome profiling experiments revealed that elevated *psm⍺* transcript levels in the *ΔrsaE* strain led to increased ribosome recruitment to *psm⍺1*, *psm⍺2*, and *psm⍺4* coding sequences and ribosome-binding regions. This was not unexpected given the higher transcript abundance. However, *psm⍺3* was a notable exception: despite increased transcript levels, ribosome occupancy remained low. SHAPE-MaP structural probing data provide an explanation: the *psm⍺3* region is highly structured, with the Shine-Dalgarno (SD) sequence sequestered within a stable stem-loop that appears to be the primary determinant limiting *psm⍺3* translation, preventing ribosome access even when transcripts are abundant. We therefore propose that the direct RsaE::*psm⍺* base-pairing (in particular the *psm⍺3* interaction) serves primarily as a fine-tuning mechanism that acts on top of the major transcriptional control exerted through *agr* and the translational constraints imposed by the RNA secondary structure.

### Deletion of RsaE activates the *agr* quorum sensing machinery

We show that the increased steady state levels of the *psm⍺* transcript levels in the *ΔrsaE* strain is likely due to (indirect) activation of the *agr* quorum sensing system, which is based on increased ribosome occupancy on *agr,* LacZ/β-galactosidase reporter assay results that showed increased activation of the *psm⍺* promoter in *ΔrsaE* and increased transcript levels of ⍺ and β toxins. Strikingly, removing RsaE did not significantly alter *agr* transcript levels, but ribosome occupancy was significantly increased. This hints that RsaE somehow influences the translation of these transcripts. We considered the possibility that RsaE directly regulates translation of mRNAs encoded within this operon. However, IntaRNA [41] and TargetRNA3 [42] did not identify promising interactions between RsaE and *agr* mRNA transcripts. Mining of existing RNA-RNA interactome data [12,14,43] also did not offer experimental evidence for direct regulation of *agr* by RsaE.

The *agr* activation in the *ΔrsaE* strain is surprising given that TCA cycle enzymes are upregulated in *ΔrsaE* and previous work established an inverse relationship between TCA cycle activity and *agr* activity in *S. aureus*. TCA cycle mutants (such as aconitase-deficient strains [44]) showed increased RNAIII levels, while forcing TCA cycle activation decreased RNAIII and altered virulence factor production [45]. Similarly, disruption of electron transport chain components resulted in decreased *agr* expression and cytolytic toxin production [46]. However, our new data, combined with previous work (reviewed in [47]), suggest that RsaE deletion leads to coordinate derepression of both TCA cycle genes and *agr* translation, which appears contrary to this established inverse relationship. This could indicate that moderate changes in TCA cycle gene expression have different regulatory outcomes than complete enzyme inactivation, that RsaE regulates these pathways independently rather than through metabolic coupling, or that the TCA-*agr* relationship is non-linear or strain-specific. Our current model is that *agr* activation by RsaE deletion is likely indirect, but the precise mechanism and its relationship to metabolic gene expression remains unclear.

### Does RsaE deletion increase alpha toxin production and secretion?

We previously showed that deleting RsaE in USA300 reduced the level of PSM⍺1 and PSM⍺4 peptides in culture supernatants and reduced the haemolytic activity [14]. Our current finding that RsaE deletion increases *psm⍺* transcript levels and ribosome occupancy on the operon is therefore unexpected, as this would normally suggest enhanced toxin production. These two independent pieces of data are not straightforward to reconcile.

However, it is possible that while more PSM⍺1 and PSM⍺4 might be produced in *ΔrsaE,* only a relatively small fraction gets secreted to culture supernatants. To address this issue, we attempted to establish absolute quantification protocols for PSM⍺ toxins using mass spectrometry, to enable direct comparison of their intracellular and extracellular levels.

However, this has proven technically very challenging, and we have not yet succeeded in developing a robust and reproducible method.

### RsaE is required for biofilm maturation, viability and dissemination

Our results provide strong evidence for an important role for RsaE in regulating biofilm maturation and thickness. We observed that *ΔrsaE* cells reproducibly produced thinner biofilms with a minimal eDNA layer on the surface. These data therefore imply an important role for RsaE in biofilm structuring and production of the protective eDNA layer in S. aureus. These results are reminiscent of the role of RsaE in *Staphylococcus epidermidis*, where RsaE promotes eDNA release and extracellular matrix production through interaction with the antiholin-encoding *lrgA* mRNA, thereby facilitating localised bacterial lysis and eDNA release into the biofilm matrix [33]. Whilst the molecular mechanism by which RsaE influences eDNA architecture in *S. aureus* remains to be fully elucidated, our findings suggest that this role in biofilm structuring may be conserved across staphylococcal species, albeit potentially through distinct target mRNAs, given that the RsaE–*lrgA* interaction was shown to be species-specific [33]. During early stages of biofilm formation *in vitro*, cell viability was significantly increased in the *ΔrsaE* strain, which may in part be linked with derepression of TCA cycle enzymes and arginine catabolism pathways [12]. This led us to hypothesise that elevated metabolic activity and cell viability might enhance bacterial dissemination from biofilms during infection. However*, in vivo* testing revealed a strikingly different outcome. Using a murine catheter model, *ΔrsaE* biofilms showed significantly reduced dissemination to kidneys after three days of infection compared to wild-type. Moreover, cryo-SEM imaging revealed that *ΔrsaE* biofilms exhibited distinct structural features *in vivo*, including sparser bacterial colonisation and gelatinous matrix production. Such sparser colonisation was not evident in laboratory culture conditions, underscoring the importance of performing *in vivo* infection studies. Longer term goals are to further dissect these biofilms grown on catheters, for example, by further characterising the protein composition of the matrix, the RNA steady state levels to look at gene expression profiles, and to analyse their resistance to attacks by the host immune system. Imaging these biofilms using CLSM to study eDNA production in the parental and mutant strains may also be informative, although protocols for performing these analyses whilst preserving the integrity of these biofilms will need to be established.

## Materials and Methods

### Bacterial strains and culture conditions

All bacterial strains and plasmids used in this study are listed in Tables 1 and 2. The *E. coli* DH5⍺ strain was used for general plasmid propagation, while the *S. aureus* RN4220 strain (SauI restriction system mutated, *hsdR* inactivated, restriction deficient) was used as an intermediate for transforming plasmids into *S. aureus* USA300 LAC. *E. coli* and *S. aureus* were cultivated in lysogeny broth (LB) and tryptic soy broth (TSB) media, respectively. Incubation was carried out at 37°C with constant shaking at 200 rpm. When necessary, selective antibiotics were incorporated into the growth media. Chloramphenicol was used at 15 μg/mL for *E. coli* and 10 μg/mL for *S. aureus*.

**Table 1.** Bacterial strains used in this study.

| Name | Purpose | References |
| --- | --- | --- |
| *E. coli* DH5α | Plasmid propagation and recombinant plasmid construction | Subcloning Efficiency™ DH5α Competent Cells (Invitrogen, 18265017) |
| *S. aureus* RN4220 | Restriction-deficient derivative of reference strain NCTC8325-4 | [48] |
| *S. aureus* RN450 80α phage | Encodes for Φ80a phage under the control of a mitomycin responsive promoter | [49] |
| *S. aureus* HG003 | Laboratory *S. aureus* strain | [50] |
| SAPhB404 | HG003 but with RsaE deleted | [51] |
| *S. aureus* USA300 LAC | USA300 clinical isolate of MRSA | [52] |
| USA300 LAC Δ*rsaE* | USA300 lacking RsaE | [14] |
| USA300 LAC Δ*rny* | USA300 lacking endo-ribonuclease RNase Y | This study |
| USA300 LAC Δ*rnc* | USA300 lacking endo-ribonuclease RNase III | This study |
| USA300 LAC:: $\Delta psm\alpha$ | Eliminate endogenous expression of *psm\alpha* operon | This study |
| USA300 LAC:: $\Delta rsaE$ + pICS3-RsaE_WT | | This study |
| USA300 LAC:: $\Delta rsaE$ + pICS3-RsaE mut-1 | | This study |
| USA300 LAC:: $\Delta rsaE$ + pICS3-RsaE mut-2 | | This study |
| USA300 LAC:: $\Delta rsaE$ + pICS3-RsaE mut-both | | This study |

**Table 2.** Plasmids used in this study: All plasmid vectors used in this study.

| Name | Purpose | Selection marker | Source |
| --- | --- | --- | --- |
| pIMAY* | Gene manipulation of *S. aureus* through allelic exchange via homologous recombination (Addgene plasmid #121441) | Cm<sup>r</sup> | [53] |
| pICS3 | *E. coli S. aureus* shuttle vector for complementing RsaE WT and mutants in USA300 LAC:: $\Delta rsaE$ strain | Amp<sup>r</sup> (*E. coli*); Cm<sup>r</sup> (*S. aureus*) | [54] |
| pICS3-RsaE mut-1 | For an *in vivo* complementation gene assay cloning RsaE with the 5' seed sequence mutated | Cm<sup>r</sup> | This study |
| pICS3-RsaE mut-2 | For an *in vivo* complementation gene assay cloning RsaE with the 3' seed sequence mutated | Cm<sup>r</sup> | This study |
| pICS3-RsaE mut-both | For an *in vivo* complementation gene assay cloning RsaE with both 5' and 3' seed sequences mutated | Cm<sup>r</sup> | This study |

### Complementation of USA300::*ΔrsaE* strain using pICS3

G-blocks containing the RsaE seed sequence mutations, along with the upstream pamiA promoter sequence, were designed (Table 3). Restriction sites were introduced between the promoter and coding sequence, as well as at both ends, to facilitate ligation into the pICS3 vector. The restriction enzymes selected were PstI and EcoRI, as both are present in the multiple cloning site (MCS) of the vector. Lyophilized Gblocks were ordered from Integrated DNA Technologies (IDT). To amplify the G-blocks, they were first cloned into the pJET1.2/blunt end cloning vector using the CloneJET kit (Thermo Scientific, K1232). After successful transformation, Positive clones were verified by Sanger sequencing. Next, G-blocks were amplified using PCR with Q5 High-Fidelity DNA Polymerase (NEB, M0491) to minimize errors during strand synthesis. The purified fragments and the pICS3 vector were then digested with PstI-HF (NEB R3140) and EcoRI-HF (NEB R3101). An overnight ligation at 16°C was set up using T4 DNA Ligase (NEB, M0202). Following heat-shock transformation in *E. coli* DH5⍺, colonies growing on LB agar plates supplemented with 10 µg/mL ampicillin were screened by colony PCR. Positive clones were grown overnight in LB with 10 µg/mL ampicillin and sent for Sanger sequencing. After verification of the recombinant plasmid sequence, it was electroporated into RN4220. Positive clones following electroporation were identified by colony PCR. Confirmed clones were then transferred into the USA300::*ΔrsaE* parent strain via phage transduction via *S. aureus* RN450 80⍺ phage. Plasmid expression was subsequently verified by northern blot using RsaE-specific probes (Table 4).

**Table 3.**
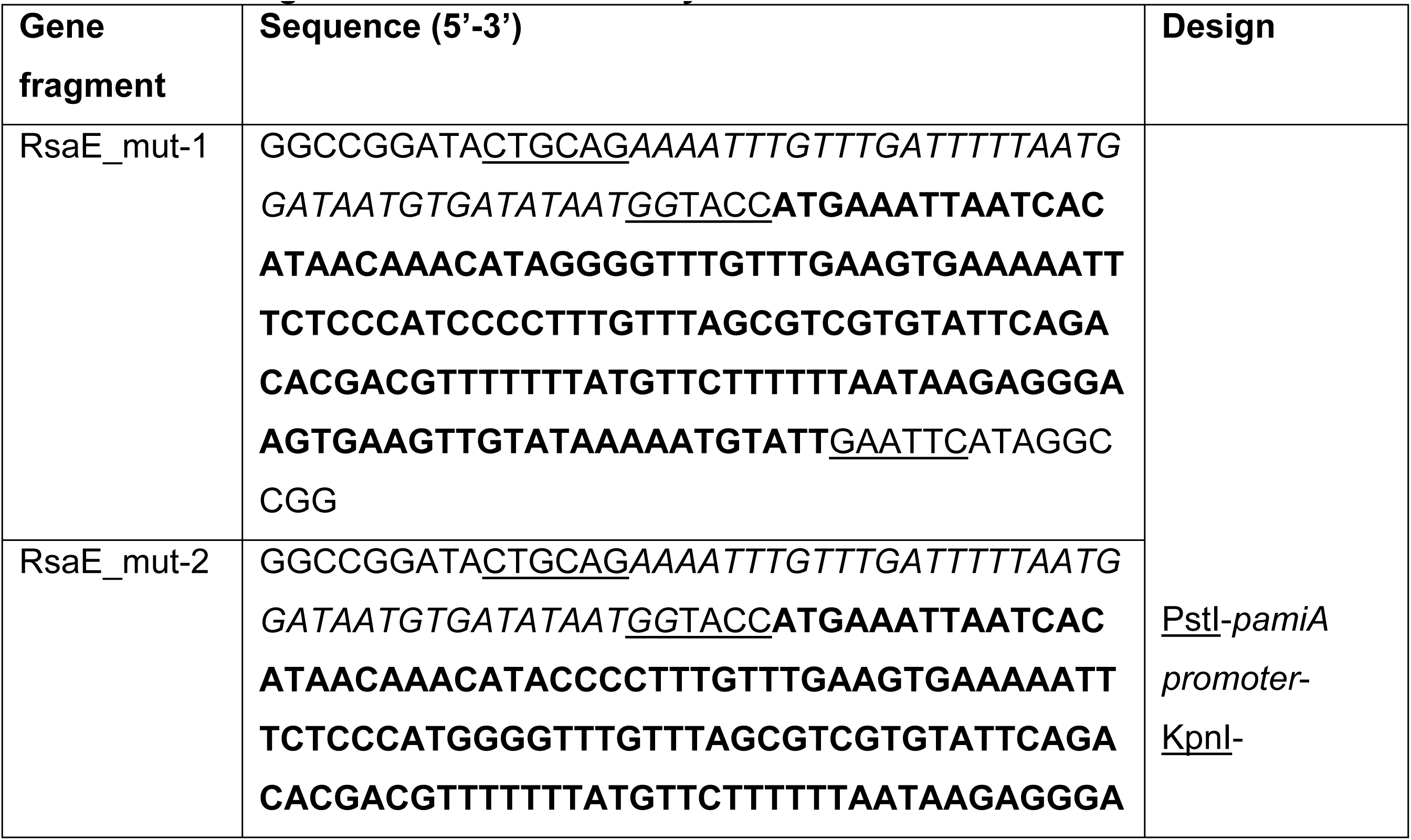
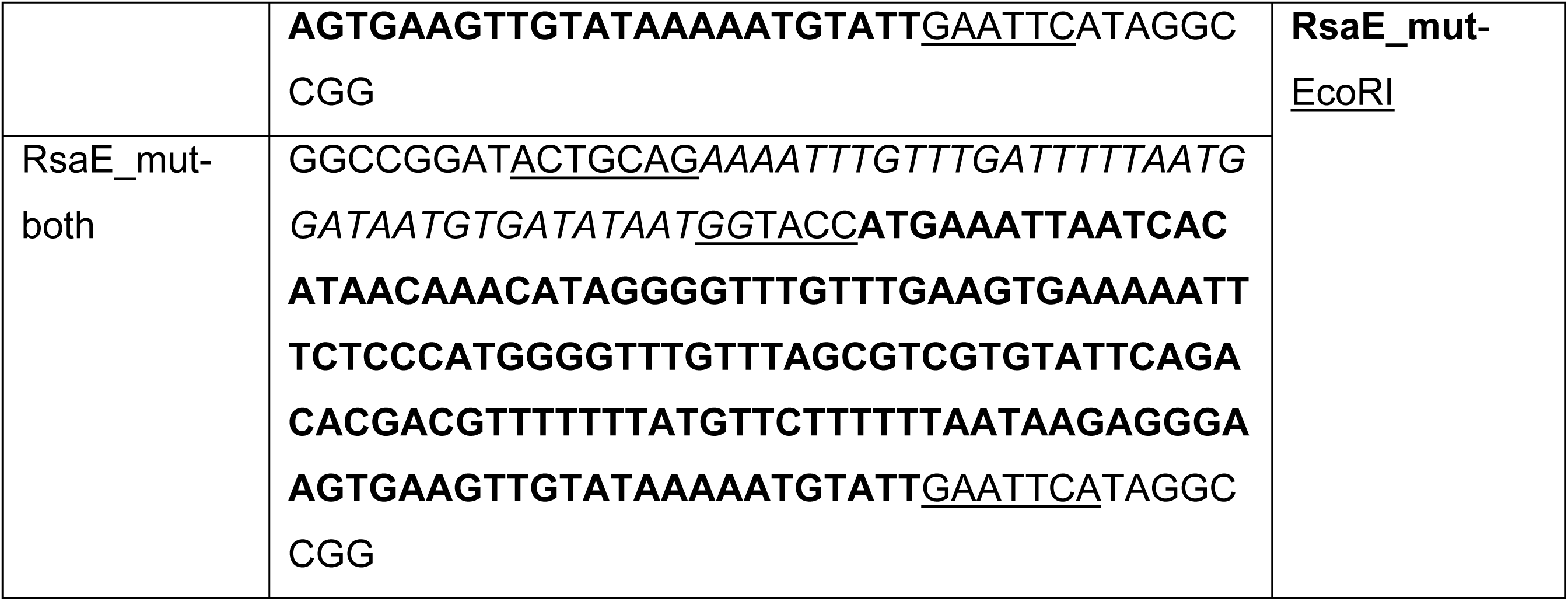
Gene fragments used in this study.

**Table 4.**
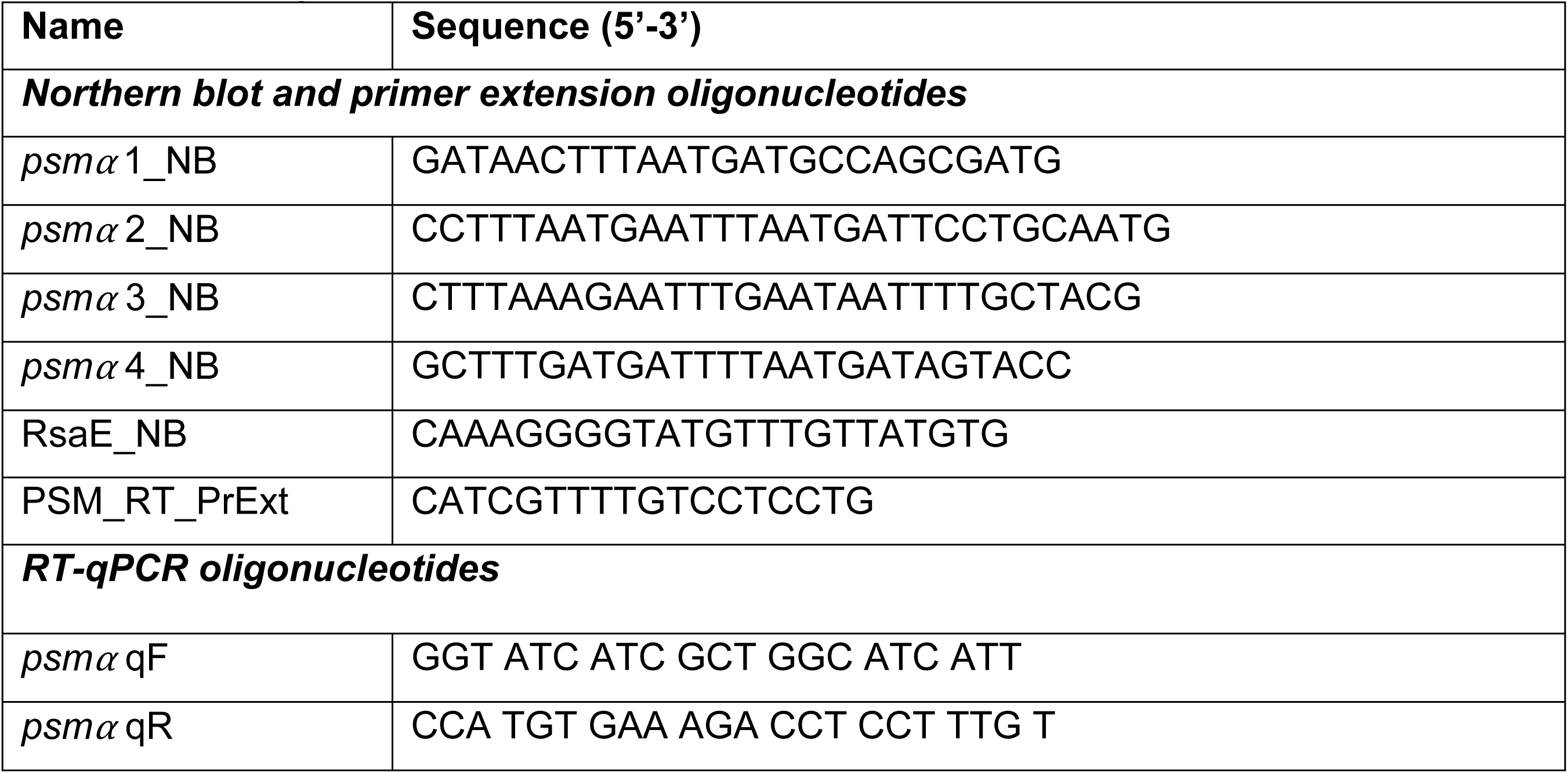
DNA oligonucleotides for Northern blot, primer extension and RT-qPCR.

### Total RNA extraction

Total RNA was extracted from strains grown to OD_595_ ∼ 3.0 using guanidium thiocyanate, acidic phenol: chloroform-based extraction [14]. Pellets were resuspended in 650 µL of guanidium thiocyanate (GTC)-phenol mix (pH 5.4; 1:1 ratio) on ice. To facilitate cell lysis, zirconia/ silica beads (diameter 0.1 mm; BioSpec products, 11079101z) were added and samples were vortexed with interspersed rests on ice. The samples were incubated at 65°C for 10 minutes and then placed on ice for 10 minutes. To each tube, 80 µL of 100 mM sodium acetate (NaOAc, pH 5.2) mix and 300 µL of chloroform:isoamyl alcohol (IAA) (24:1) were added, followed by vortexing and centrifugation at 12,045 x g for 2 minutes. The upper aqueous phase containing RNA was carefully transferred to new tubes containing 500 µL of Phenol:Chloroform:Isoamyl alcohol mix (25:24:1), vortexed vigorously, and centrifuged again at 12,045 x g for 2 min This step was repeated once, after which the aqueous phase was transferred to tubes containing 500 µL of chloroform:IAA (24:1) and centrifuged again to complete phase separation. Following the final phase separation, ∼500 µL of the aqueous phase was transferred to tubes containing 2.5 volumes of ice-cold 96% ethanol and 20 µg of glycogen for RNA precipitation. After a thorough vortex samples were stored at -80°C for at least 30 min or at -20°C overnight. Samples were then centrifuged at 12,045 x g for 30 min at 4°C, the supernatant was removed, and the pellets were washed with 1 mL of ice-cold 70% ethanol. After another centrifugation at 12,045 x g for 5 min, the ethanol was removed completely, and pellets were air-dried. Finally, RNA pellets were resuspended in DEPC-treated water (Invitrogen, AM9922).

RNA concentration was measured using Qubit RNA HS assay kits (Invitrogen, Q32852) in Qubit 4 systems and RNA integrity was checked using Agilent RNA 6000 Nano kit (Agilent, 5067-1511) with Bioanalyzer 2100 (Agilent Technologies).

### Northern Blot analysis

Six percent PAA-urea gel (6 M urea, 1X TBE, 6% (v/v) acrylamide, 0.05% APS, and 0.05% TEMED) was casted between 2 mm thick, 20x20 cm Notched glass plates (Fisher Scientific, 1185453) using a 25 mL serological pipette. The gel thickness was set using 1 mm bonded spacers (Fisher Scientific). Five µg of RNA samples and RNA Century Marker Plus ladder (Invitrogen, AM7145) mixed with 2X RNA loading dye (Thermo Scientific, R0641) were denatured at 85°C for 3 minutes followed by incubation on ice for 5 minutes. Samples were resolved at 500 V in 1X TBE. For Northern blot analyses, RNA in the gel was electroblotted to a nitrocellulose membrane (Amersham Hybond-N^+^, RPN203B) at 50 V for 4 hours, then crosslinked using 1200 mJ of 254 nm UV radiation. The membrane was prehybridized in 7 mL of UltraHyb Oligo buffer (Invitrogen, AM8663) at 42°C for 1 hour. A radiolabelled DNA probe was then added to the prehybridization solution for overnight incubation at 42°C. After hybridization, the membrane was washed twice with 2X SSC (0.03M Sodium Citrate, 0.3 M NaCl), 0.5% SDS. Imaging was performed with a Typhoon™ FLA 7000 biomolecular imager (GE Healthcare) using radiosensitive imaging screen (GE Healthcare).

### Rifampicin mRNA stability assay

To grow the cells, fresh cultures were inoculated at an OD_595_ 0.05 in 250 mL flasks ensuring that the media-to-flask ratio was approximately 1:6. The cultures were grown overnight at 37°C with shaking at 180 rpm. Once the culture reached mid-exponential phase (OD_595_ 0.4-1), a 5 mL sample was taken at time point 0 sample, and the cells were pelleted. Next, Rifampicin (Sigma Aldrich, R3501) was added to the culture at a final concentration of 200 µg/mL. At each subsequent time point (5, 15, 30, and 60 minutes), 5 mL samples of the culture were collected, and the cells were pelleted. RNA extraction was then performed on these samples and RNA was resolved on 8% 7M Urea polyacrylamide gels.

### Reverse Transcription-quantitative PCR (RT-qPCR)

For each RNA sample, DNase treatment was performed with 4 U RQ1 DNase (Promega, M6101) according to manufacturer’s guidelines. 2 U SUPERase·In™ RNase Inhibitor was also included in the mix. The RNA was then purified using RNAClean XP beads (Beckman Coulter). Reverse transcription was carried out using SuperScript™ IV (SS IV) Reverse Transcriptase (Invitrogen, 18090010). The qPCR master mix was prepared by combining 1X Brilliant III Ultra-Fast SYBR® Green QPCR Master Mix (Agilent, 600883) with 4 µM of the primer pair mix. The qPCR was performed in Lightcycler 480 (Roche). Relative gene expression levels were determined using the 2×ΔΔCt method, normalizing against 5S rRNA as an internal reference. Each reaction was performed in technical triplicate, and data represent the mean and standard error of mean (SEM) of three independent biological replicates.

### USA300 RNA-seq experiments

USA300 WT and *ΔrsaE* strains were grown in TSB to an OD_595_ ∼ 3.0, starting from an OD_595_ of 0.05. Five mL of culture was harvested and pelleted by centrifugation at 5,000 × g for 5 minutes. RNA was extracted from USA300 LAC strains as previously described [14]. rRNA-depleted RNA was sequenced by SeqCentre using the TruSeq protocol.

### *In vivo* structure probing of total RNA

Cells were grown to OD_595_ ∼ 3.0 in TSB, starting from an OD_595_ of 0.05. Five mL of culture was harvested and pelleted by centrifugation at 5,000 × g for 5 minutes. The pellet was resuspended in 450 µL of 1X PBS. In parallel, 50 µL of 1M 2A3 ((2-Aminopyridin-3-yl) (1H-imidazol-1-yl) methanone) was thawed and vortexed extensively before being added to the cell suspension to achieve a final concentration of 100 mM 2A3. For the vehicle control, 50 µL of DMSO was added instead. The suspension was mixed by pipetting and incubated at 37°C for 25 min with shaking. Following incubation, the reaction was quenched by adding 500 mM Dithiothreitol (DTT) (Thermo Scientific, R0862). The cells were then pelleted by centrifugation at 4°C and taken for RNA extraction. RNA was extracted followed by DNase treatment with 4 U RQ1 DNase (Promega, M6101) and 2 U SUPERase·In™ RNase Inhibitor (Invitrogen, AM2696) at 37°C for 1 hour. RNA was purified using RNAClean XP beads.

### SHAPE-MaP library preparation

One µg of DNase treated RNA was taken up for Ribosomal RNA (rRNA) depletion using NEBNext rRNA depletion kit for Bacteria (NEB, E 7850). rRNA depleted RNA was then fragmented to an average length of 150 nucleotides by incubating at 94°C for 8 minutes in a fragmentation buffer (65 mM Tris-HCl (pH 8.0), 95 mM KCl, and 4 mM MgCl_2_). Following fragmentation, RNA was purified using RNAClean XP beads. Reverse transcription was performed using SuperScript™ II Reverse Transcriptase (Invitrogen, 18064014). The reaction mixture contained 5 µL of fragmented RNA, 0.4 µL of 50 µM random hexamers (Invitrogen, N8080127) and 0.5 µL of 10 mM dNTPs. The mixture was incubated at 70°C for 5 minutes, then placed on ice for 1 minute to facilitate primer annealing. A reverse transcription master mix was then prepared, consisting of 1X First Strand (FS) Reaction Buffer, 0.01 M DTT, 6 mM MnCl_2_, 5 U SUPERase•In™ RNase Inhibitor, and 50 U SSII, with the final reaction volume adjusted to 10 µL. The reaction was incubated at 25°C for 10 minutes to allow partial primer extension, followed by a 3-hour incubation at 42°C for cDNA synthesis. SSII was heat-inactivated at 72°C for 10 minutes. To remove Mn^2+^ ions, 6 mM final EDTA was added, followed by incubation at room temperature for 5 minutes. To restore Mg^2+^ concentrations required for the subsequent steps, 6 mM final MgCl_2_ was added. Second strand synthesis was performed using the NEBNext® ultra II Non-Directional RNA Second Strand Synthesis Module (NEB, E6111S), following the manufacturer’s instructions. The reaction was incubated at 16°C for 1 hour. cDNA libraries were then prepared using the NEBNext® ultra™ II DNA Library Prep Kit for Illumina® (NEB, E7645S), according to the manufacturer’s protocol. The library prepared was sent for RNAseq.

### *In vitro* biofilm analysis

*S. aureus* cultures were streaked on TSA plates and incubated overnight at 37°C. A single colony was then inoculated into 5 mL of TSB and grown overnight at 37°C with shaking. To synchronise the cultures, fresh cultures were initiated at an OD_595_ of 0.05 in fresh TSB. Subsequently, 10 μL of culture at OD_595_ 0.5 was inoculated into 190 μL of TSB supplemented with 0.5% glucose in 96-well microtiter plates. For cultures intended for confocal imaging, clear-bottom black 96-well plates from ibidi® (ibiTreat µ-Plate, Cat. No. 89626) with a polymer coverslip designed for imaging were used. For all other applications, Corning® Costar® tissue culture-treated transparent 96-well plates (Cat. No. CLS3516) were used. Plates were incubated statically at 37°C for 24, 48, and 72 hours.

### *In vivo* biofilm analysis

1. *S. aureus* cultures were streaked on TSA plates and incubated overnight at 37°C. A single colony was then inoculated into 5 mL of TSB and grown overnight at 37°C with shaking.

To synchronise the cultures, fresh cultures were initiated at an OD_595_ of 0.05 in fresh TSB. Subsequently, 10 μL of culture at OD_595_ 0.5 was inoculated into 190 µL of TSB supplemented with 0.5% glucose in 24 well tissue culture plates (TPP, R2012) containing 1 cm catheters (Durect Corporations, Cat. No. NC9166951) cut into halves which had been treated overnight with human plasma (3.2% NaCit, 4 individuals mixed gender pooled, 0.22 µm filtered) to facilitate bacterial adherence. After 24 hours, the media was aspirated and catheters were washed in 1X PBS to remove planktonic cells before insertion into mice.

### Mice infection

Murine work was carried out according to protocol #00000081 approved by the Ohio University Institutional Animal Care and Use Committee (IACUC). On the day of infection, mice were anesthetised with isoflurane. Immediately following anaesthetic induction, ophthalmic ointment (OptixCare, 77113) was applied to the animals’ eyes to prevent corneal drying. Once anesthetized, a small patch of fur (approximately 2 cm x 2 cm) was shaved on the dorsum. The surgical site was scrubbed beginning at the centre and moving outward in concentric circles using either dilute chlorhexidine or povidone iodine scrub, followed by 70% isopropyl alcohol or sterile saline. A local analgesic, 0.25% Bupivacaine Hydrochloride injection (2.5 mg/mL, 0.1-0.2 mL) was injected subcutaneously at the surgical site. An approximately 1 cm incision was made in the centre of the shaved area. Two sections of catheter tubing were then implanted subcutaneously in each mouse, one to the left and one to the right of the incision. A haemostat was used if necessary to facilitate catheter placement. After implantation, the incision was closed using sterile 7 mm wound clips. The mice were euthanised at Day 3 of infection and catheters were removed for performing CFU count and SEM.

### Biomass estimation of *in vitro S. aureus* biofilms

After incubation, non-adherent cells and media were removed by pipetting. 3.7% formaldehyde was used to fix the biofilm. A 0.1% Crystal Voilet solution was added to each well and incubated at room temperature for 15 minutes. Wells were subsequently rinsed three times with 1X PBS to remove excess dye. The bound CV was solubilised using 30% acetic acid, and absorbance was measured at 575 nm using 30% acetic acid as the blank.

### CFU analysis of *in vitro* biofilms

After biofilms were grown for the desired time points, non-adherent cells and media were removed from each well by pipetting. Wells were washed with 200 µL of 1X PBS to eliminate remaining planktonic cells. To prevent contamination, the wells were covered with parafilm, and the plate lid was double-sealed. Plates were then sonicated in a water bath (Branson 2510) for 5 minutes to dislodge biofilm-associated cells. Following sonication, the released cells were resuspended by pipetting to ensure uniform distribution. A 10^-8^ serial dilution was prepared, and 50 µL of each dilution was plated onto TSA or TSA supplemented with appropriate antibiotics, depending on the bacterial strain. Two-compartment 94/15 mm Petri dishes (Greiner Bio-One, Cat. No. 635161) were used. Plates were incubated overnight, and colony-forming units (CFU) were counted the following day using OpenCFU software [55] to determine CFU/mL.

### CFU analysis of *in vivo* biofilms

Kidney samples were homogenised using a Mini-Beadbeater 16 (Biospec Products) with 2.3 mm zirconia/silica beads (Biospec Products, 11079125Z), and 100 µL of 1X Phosphate Buffered Saline (PBS) to ensure thorough mechanical disruption and homogenisation of the tissue material. 10^-1^ to 10^-8^ dilution series were prepared. 50 µL of the diluted samples were then plated onto Manitol salt agar (MSA) to select for *Staphylococcus aureus*. The plates were incubated overnight, and the colonies were counted the next day using OpenCFU software [55] to determine CFU/mL.

### Confocal microscopy for extracellular polymeric substances (EPS) distribution analysis in *in vitro* biofilms

The biofilms were grown up to desired time point. Following removal of planktonic cells, 3.7% formaldehyde was added to the wells and incubated for 30 minutes at room temperature. Excess formaldehyde was removed by washing the wells with 1X phosphate-buffered saline (PBS). FilmTracer™ SYPRO™ Ruby Biofilm Matrix Stain (Invitrogen, F10318) was used to fluorescently stain proteins, Wheat Germ Agglutinin (WGA) Alexa Fluor 488 (Invitrogen, W11261) was used to stain polysaccharides, while Hoechst 33342, trihydrochloride trihydrate (Invitrogen, H3570) was used to stain nucleic acids, i.e., bacteria within the EPS. After fixation, 200 μL of 1X SYPRO Ruby dye was added to the wells containing samples, as well as to an empty well serving as a control. The plate was covered with aluminium foil to prevent light exposure and incubated at room temperature (RT) for 15 minutes. Following incubation, three consecutive washes with autoclaved distilled water were performed. PBS was not used for washing, as phosphate ions interfere with dye binding according to the manufacturer’s instructions. Next, 200 μL of a staining solution containing 5 μg/mL Hoechst and 5 μg/mL WGA in autoclaved distilled water was added to the wells. After incubation at RT for 15 minutes whilst covered with aluminium foil, three consecutive washes with autoclaved distilled water were performed.

For eDNA visualization, TOTO-3 iodide (Invitrogen, T3604) was used to stain B-DNA and Z22 antibody, Anti-Z-DNA/Z-RNA [Z22] (Absolute Antibody, Ab00783-23.0) was used to stain Z-DNA, while Hoechst 33342, trihydrochloride trihydrate (Invitrogen, H3570) was used to stain nucleic acids, i.e., bacteria within the EPS. Since the Z-22 antibody did not have a probe, an Alexa Fluor™ Plus 594 conjugated Donkey anti-Rabbit IgG (H+L) ((Invitrogen, A32754) was used as secondary. Following fixation, 100 μL of 3% bovine serum albumin (BSA) in 1X PBS was added to each well to reduce nonspecific binding. After a short blocking step, 100 μL of Z-22 antibody diluted 1:100 in 3% BSA was added and incubated for 1 hour at RT. Wells were then washed with 100 μL of 3% BSA in 1X PBS. A fluorescently labelled secondary antibody, Donkey anti-Rabbit IgG (H+L) Alexa Fluor™ Plus 594, diluted 1:150 in 3% BSA, was added and incubated for 1 hour at RT. Excess antibody was removed by washing with 3% BSA. As a final step, 20 μg/mL Hoechst and 2 μM TOTO-3 were added and incubated for 30 minutes before washing with 1X PBS. The stained biofilms were then visualised using a Nikon Ti2 Spinning Disk Confocal Microscope with the following excitation and emission filters: 450 nm excitation / 610 nm emission for SYPRO Ruby, 495 nm excitation / 519 nm emission for WGA, 350 nm excitation / 461 nm emission for Hoechst, 642 nm / 660 nm for TOTO-3 and 561 nm / 630 nm for Alexa Fluor 594.

### SEM sample preparation of *in vivo* biofilms

Catheters with biofilm formation were aseptically retrieved from mice and immediately fixed in 2.5% glutaraldehyde solution at 4°C overnight. Following primary fixation, the samples were further fixed in increasing concentration of Ethanol, 15%, 30%, 50%, 70%, 90%, and 100%, with each step carried out for 15 minutes at RT. Dehydrated samples were subsequently dried using a vacuum centrifuge for 5 minutes. The catheters were then mounted onto aluminium stubs using double-sided adhesive tape and sputter-coated with gold prior to imaging.

### SEM imaging of biofilms

We used a Hitachi S-4500FE SEM (10 uA current and 20 kV accelerating voltage), Oxford AZtec software to capture images, and a Pelco SC-4 to sputter the gold coating on the samples (operated at 10 mA, typically sputters 3 Angstroms/second). After mounting the samples on aluminum stubs using carbon tape, a ∼40 nm layer of Au was applied to the surface of the catheters using a Pelco SC-4 sputter coater. The samples were then transferred to the Hitachi S-4500FE SEM where representative clusters of cells were located and imaged at magnifications of 3,000X and 1,500X for each catheter. Images were captured using Oxford Instrument’s AZtec software.

### Ribosome profiling analysis

HG003 and HG003 *ΔrsaE S. aureus* overnight cultures were grown in Brain Heart Infusion (BHI) broth at 37°C with agitation. Cultures were diluted 100-fold into fresh BHI medium and grown to an optical density at 600 nm (OD₆₀₀) of 1.0. Cultures were treated with chloramphenicol (100 µg/mL) for 2 min and then rapidly cooled on ice and harvested by centrifugation at 4700 rpm at 4°C. Pellets were flash-frozen in liquid nitrogen and stored at −80°C until further use. Cell pellets were resuspended in 200 µL lysis buffer (20 mM Tris-HCl pH 8.0, 100 mM NH2Cl, 150 mM magnesium acetate, 0.4% Triton X-100, 0.2% NP-40, 2 × EDTA-free protease inhibitor cocktail (Roche), 60 Units murine RNase inhibitor, 2 Units Turbo DNase I per 50 µg RNA (Promega). Lysis was performed using a FastPrep-24 instrument (MP Biomedicals) with glass beads (0.1 mm diameter), applying three cycles of 42 s shaking followed by 5 min cooling on ice. Lysates were clarified by centrifugation at 21,000 × g for 5 min at 4°C, and supernatants were stored at −80°C. An aliquot (∼50 µL) was retained for total RNA extraction. Lysate RNA content was estimated by measuring absorbance at 260 nm using a NanoDrop (Thermo Fisher Scientific) spectrophotometer. Background from lysis buffer was subtracted.

Polysomes were pelleted through a 2 mL 30% (w/v) sucrose cushion (50 mM Tris-acetate pH 7.6, 50 mM NH4Cl, 12 mM MgCl2, 1 mM DTT) by ultracentrifugation in a TLA110 rotor (Beckman Coulter) at 100,000 rpm for 90 min at 4°C. Pellets were washed in polysome resuspension buffer (20 mM Tris-HCl, pH 8.0, 10 mM MgCl2, 100 mM NH4Cl, 5 mM CaCl2, 1× BSA) and resuspended in 250 µL of the same buffer. Micrococcal nuclease (MNase; New England Biolabs) was added at 100 Units per A_260_ unit and incubated at 25°C for 1 h. Digestion was terminated by addition of 7.5 mM EGTA (pH 8.0). To evaluate digestion efficiency, selected samples were loaded onto 31% sucrose gradients prepared in 50 mM Tris-acetate (pH 7.6), 50 mM NH4Cl, 12 mM MgCl_2_, 1 mM DTT. Gradients were generated via freeze-thaw cycling at –20°C. Samples were centrifuged in an SW41 rotor at 240,000 x g at 4°C for 3 h and fractionated using a Teledyne ISCO gradient fractionator. Monosome fractions from digested samples were additionally purified via 30% sucrose cushions using the above ultracentrifugation conditions.

RNA Extraction and Size Selection: RNA was extracted from monosome fractions with 500 µL acidic phenol (pH <5.2) at 65°C for 15 min by vortexing. Following two rounds of chloroform extraction, RNA was precipitated using isopropanol and 0.3 M sodium acetate (pH 5.2), centrifuged at 21,000 × g for 20 min at 4°C, and resuspended in 100 µL RNase-free water. Concentration was determined via A260. A total of 15–32 µg RNA was resolved on a 17% denaturing polyacrylamide gel (19:1 acrylamide:bisacrylamide, 7 M urea, 1× TAE) at 100 V for 5 h. Gels were pre-run for 30 min and stained with SYBR Gold (10 µg/mL, Thermo Fisher Scientific). Fragments corresponding to 28–34 nt were excised and eluted overnight at 4°C in 0.3 M sodium acetate (pH 5.5), 1 mM EDTA with rotation. RNA was recovered through 0.45 µm Spin-X cellulose acetate columns (Costar), precipitated with ethanol and glycogen (20 mg/mL), and pelleted by centrifugation at 21,000 × g for 30 min at 4°C. Final RNA was resuspended in 15 µL RNase-free water and quantified using the Quant-iT microRNA Assay Kit (Thermo Fisher Scientific). Ribosomal RNA was depleted using the RiboPOOL kit for S. aureus (siTOOLs Biotech) according to the manufacturer’s instructions. RNA was purified using the RNA Clean & Concentrator-5 kit (Zymo Research) with additional ethanol steps to retain small RNA. For 3′ end repair, 8 µL RNA was incubated with 2 µL 10× Antarctic phosphatase buffer, 1 µL Antarctic phosphatase (New England Biolabs), 1 µL SUPERase•In (, and 8 µL water at 37°C for 30 min, followed by inactivation at 80°C for 2 min. The 5′ phosphorylation of RNA was performed in 30 µL reactions with 3 µL 10× T4 PNK buffer, 5 µL 10 mM ATP, 2 µL T4 polynucleotide kinase (New England Biolabs), 1 µL SUPERase•In, and 19 µL water at 37°C for 1 h. Reactions were heat-inactivated at 65°C for 20 min. RNA was re-purified using the RNA Clean & Concentrator-5 kit and eluted in 8 µL water.

Libraries were constructed using the NEBNext Multiplex Small RNA Library Prep Set (New England Biolabs) following the manufacturer’s instructions, using half-volume reactions and 12 PCR cycles. PCR products were purified with the Monarch PCR & DNA Cleanup Kit (New England Biolabs) and eluted in 27.5 µL. Library quality and concentration were assessed using a Bioanalyzer 2100 with High Sensitivity DNA chips (Agilent) and the Quant-iT dsDNA HS Assay (Thermo Fisher). Where necessary, secondary library products were removed by PAGE. PCR products (25 µL) were resolved on a 6% native TBE gel run at 200 V for 30 min. Gels were stained with SYBR Gold, and bands of ∼150 nt were excised. DNA was eluted overnight in 250 µL DNA elution buffer, filtered through Costar gel filtration columns, and precipitated with 25 µL 3 M sodium acetate (pH 5.5), 2 µL glycogen (10 mg/mL), and 750 µL ethanol at –80°C for ≥1 h. DNA was recovered by centrifugation (21,000 × g, 30 min, 4°C), resuspended in 10 µL RNase-free water, and quantified.

### Bioinformatics analysis CLMS image analysis

We quantified cell fluorescence using a Cellpose [35] segmentation-based Python pipeline developed by our group to quantify cell counts and mean signal intensities. All unprocessed .nd2 image files, the code to use to analyse the data and the resulting output files can be found on the Granneman lab GitLab repository (https://git.ecdf.ed.ac.uk/sgrannem/s_aureus_biofilm_analyses). The segmentation channel was deconvolved using the Richardson-Lucy algorithm (10 iterations) to enhance contrast. In-focus Z-slices were selected using a slice-wise signal-to-noise ratio metric - defined as (98th percentile - median) / median absolute deviation of the high-pass residual - and a relative-increase percentile filter (60th percentile) to capture the contiguous slice-range containing the ring signal while minimising defocus noise (minimum window = 2 slices). Prior to segmentation, channels underwent Gaussian background subtraction (σ = 4.0 µm) to reduce extracellular diffuse signal; for DAPI/Hoechst channels in the protein dataset an additional white top-hat filter (radius = 1.5 µm) was applied to isolate intracellular nucleoid signal. Three-dimensional instance segmentation was performed with Cellpose (cyto2 model) in 2D+stitch mode, in which 2D segmentation masks from consecutive Z-slices are stitched across Z by IoU-based label propagation (stitch_threshold = 0.1). Cell diameter was auto-detected per image. For each segmented object, an annular (ring) mask was constructed by EDT-based morphological dilation of the segmentation label (dilation radius = 3 pixels) and subtraction of the cell body mask. Within each ring mask we measured mean, median and integrated intensity and estimated local background from an outer annulus; ring SNR was computed as (mean_ring − median_background) / std_background. Per-cell measurements (centroid coordinates, volume, XY aspect ratio, ring mean, ring integrated intensity, ring SNR and QC flags) were saved to “*_summary_3d.csv” files. Objects with volumes outside expected ranges or implausible XY aspect ratios were excluded from downstream analyses. Key parameters used in the analysis include “flow_threshold” ≈ 0.2-0.4 and “cellprob_threshold” = 0.0-1.5 for Cellpose detection (higher cellprob_threshold applied to DAPI channels to suppress low-confidence detections arising from extracellular dye diffusion); “min_volume_um3” = 0.50 µm^3^ and “max_volume” = 1.5 µm^3^ as physical-unit volume bounds; and “max_aspect_ratio” = 3.0 to reject elongated debris and merged cell chains. Focus selection settings used “focus_percentile” = 60 and “focus_fit_threshold_fraction” = 0.5. Deconvolution, focus selection, segmentation thresholds and morphological parameters are recorded with outputs to ensure reproducibility of the data. Outputs produced per sample include deconvolved segmentation TIFF (”*_seg_channel_deconvolved.tif”), per-cell per-channel summary CSV (”*_summary_3d.csv”), visualisation overlays (Z-slice overlays with segmentation and ring), and PDF files showing biofilm thickness, cell counts, 3D rendering of the cells and centroids, the cell counts and cell mean signal intensities. All processing steps, parameter names and default values are implemented in “run_cell_counting_analyses_cellpose.py” in our code.

### SHAPE-MaP data analyses

Analysis for the SHAPE-Maq data set was performed using the RNA Framework pipeline [29]. Reads were pre-processed by trimming TruSeq adapters and 5’ homopolymers from FASTQ files. Paired-end reads from 2A3 treated and untreated samples were then processed and mapped to *S. aureus* USA300 reference genome using *rf-map* with bowtie 2 and cutadapt 5.0. Mutations were then counted using *rf-count* and SHAPE reactivities were normalised using a scoring method defined by Siegfried *et al.* [56]. The code used for analysing the SHAPE-MaP data with associated documentation is available from NCBI GEO (GSE321596).

### Ribosome profiling data analyses

Raw fastq files were mapped to the USA300 FPR3757 genome using Bowtie2 [57] to enable direct comparison with RNA-seq data generated in this strain. BAM files were subsequently further processed using the pyCRAC package (version 1.5.2; [58]) to count the number of reads that mapped to each annotated gene using our USA300 genome annotation GTF file [14]. To detect reads mapping to individual *psm⍺* toxins, we included genome coordinates from 5 nucleotides upstream of the RBS until the stop codon into the GTF file. Differential read count analyses was performed using DESeq2 [59] using default settings.

### RNA-seq data analysis

Raw fastq files were mapped to the USA300 FPR3757 genome using Bowtie2 [57] and BAM files were subsequently further processed using the pyCRAC package (version 1.5.2; [58]) to count the number of reads that mapped to each annotated gene using our USA300 genome annotation GTF file [14]. To detect reads mapping to the operon transcript, we used Differential read count analyses was performed using DESeq2 [59] using default settings.

## Data availability

All raw and processed Illumina sequencing data has been made available on NCBI Gene Expression Omnibus under the accession numbers GSE320361, GSE320367 and GSE321596. The code for analysing the SHAPE-MaP data has been made available as jupyter notebook and html file under accession number GSE321596. The pyCRAC package is available from the Granneman lab git repository (git.ecdf.ed.ac.uk/sgrannem). The CLMS imaging data, the code used to analyse the data and all the resulting output files used for generating this manuscript are available from the Granneman lab gitlab repository (https://git.ecdf.ed.ac.uk/sgrannem/s_aureus_biofilm_analyses).

## Acknowledgements

We thank Myles X. Fox for assistance preparing samples for SEM analysis, Florence Lorieux for assisting with the ribosome profiling experiments and the Granneman lab members for critically reading the manuscript. This work was supported by a Medical Research Council Programme grant (MR/Y013131/1) to S.G., a Darwin Trust PhD training fellowship to M.C., an ANR grant (ANR-19-CE12-0006) to P.B., an National Institute of Allergy and Infectious

Diseases (NIAID) grant (R01AI143743) to R.K.C and a RESEARCH INCENTIVE 12322-13-4100001-100070 grant to R.K.C.

## Author Contributions

Conceptualisation: M.C, S.G, P.B, R.K.C and S.G. Data Curation: S.G. Formal Analysis: M.C. J.R.T and S.G. Funding Acquisition: P.B, R.K.C and S.G. Investigation: All authors. Methodology: M.C, R.W.S, P.A, I.H, O.N, P.B, R.K.C and S.G. Project Administration: P.B., R.K.C. and S.G. Resources: P.B., R.K.C and S.G. Software. M.C. and S.G. Supervision: P.B, R.K.C and S.G. Validation: All authors. Visualisation: M.C. and S.G. Writing - Original Draft Preparation: M.C. and S.G. Writing - Review & Editing: All authors.

## References

1. Russell CD, Fairfield CJ, Drake TM, Turtle L, Seaton RA, Wootton DG, et al. Co-infections, secondary infections, and antimicrobial use in patients hospitalised with COVID-19 during the first pandemic wave from the ISARIC WHO CCP-UK study: a multicentre, prospective cohort study. Lancet Microbe. 2021;2: e354–e365. doi:10.1016/S2666-5247(21)00090-2

2. Naghavi M, Vollset SE, Ikuta KS, Swetschinski LR, Gray AP, Wool EE, et al. Global burden of bacterial antimicrobial resistance 1990–2021: a systematic analysis with forecasts to 2050. The Lancet. 2024;404: 1199–1226. doi:10.1016/S0140-6736(24)01867-1

3. Jenul C, Horswill AR. Regulation of *Staphylococcus aureus* Virulence. Microbiol Spectr. 2019;7. doi:10.1128/microbiolspec.GPP3-0031-2018

4. Holmqvist E, Wagner EGH. Impact of bacterial sRNAs in stress responses. Biochem Soc Trans. 2017;45: 1203–1212. doi:10.1042/BST20160363

5. Felden B, Vandenesch F, Bouloc P, Romby P. The Staphylococcus aureus RNome and Its Commitment to Virulence. PLoS Pathog. 2011;7: e1002006. doi:10.1371/journal.ppat.1002006

6. Durand S, Braun F, Helfer A-C, Romby P, Condon C. sRNA-mediated activation of gene expression by inhibition of 5’-3’ exonucleolytic mRNA degradation. Elife. 2017;6. doi:10.7554/eLife.23602

7. Durand S, Tomasini A, Braun F, Condon C, Romby P. sRNA and mRNA turnover in Gram-positive bacteria. FEMS Microbiol Rev. 2015;39: 316–330. doi:10.1093/femsre/fuv007

8. Bonnin RA, Bouloc P. RNA degradation in staphylococcus aureus: Diversity of ribonucleases and their impact. International Journal of Genomics. Hindawi Publishing Corporation; 2015. doi:10.1155/2015/395753

9. Bronesky D, Wu Z, Marzi S, Walter P, Geissmann T, Moreau K, et al. *Staphylococcus aureus* RNAIII and Its Regulon Link Quorum Sensing, Stress Responses, Metabolic Adaptation, and Regulation of Virulence Gene Expression. Annu Rev Microbiol. 2016;70: 299–316. doi:10.1146/annurev-micro-102215-095708

10. Geissmann T, Chevalier C, Cros M-J, Boisset S, Fechter P, Noirot C, et al. A search for small noncoding RNAs in Staphylococcus aureus reveals a conserved sequence motif for regulation. Nucleic Acids Res. 2009;37: 7239–7257. doi:10.1093/nar/gkp668

11. Sorensen HM, Keogh RA, Wittekind MA, Caillet AR, Wiemels RE, Laner EA, et al. Reading between the Lines: Utilizing RNA-Seq Data for Global Analysis of sRNAs in Staphylococcus aureus. mSphere. 2020;5. doi:10.1128/MSPHERE.00439-20

12. Rochat T, Bohn C, Morvan C, Le Lam TN, Razvi F, Pain A, et al. The conserved regulatory RNA RsaE down-regulates the arginine degradation pathway in Staphylococcus aureus. Nucleic Acids Res. 2018;46: 8803–8816. doi:10.1093/nar/gky584

13. Bohn C, Rigoulay C, Chabelskaya S, Sharma CM, Marchais A, Skorski P, et al. Experimental discovery of small RNAs in Staphylococcus aureus reveals a riboregulator of central metabolism. Nucleic Acids Res. 2010;38: 6620–6636. doi:10.1093/nar/gkq462

14. McKellar SW, Ivanova I, Arede P, Zapf RL, Mercier N, Chu L-C, et al. RNase III CLASH in MRSA uncovers sRNA regulatory networks coupling metabolism to toxin expression. Nat Commun. 2022;13. doi:10.1038/s41467-022-31173-y

15. Otto M. Phenol-soluble modulins. International Journal of Medical Microbiology. 2014;304: 164–169. doi:10.1016/j.ijmm.2013.11.019

16. Dastgheyb SS, Villaruz AE, Le KY, Tan VY, Duong AC, Chatterjee SS, et al. Role of Phenol-Soluble Modulins in Formation of Staphylococcus aureus Biofilms in Synovial Fluid. Infect Immun. 2015;83: 2966–2975. doi:10.1128/IAI.00394-15

17. Joo H-S, Cheung GYC, Otto M. Antimicrobial Activity of Community-associated Methicillin-resistant Staphylococcus aureus Is Caused by Phenol-soluble Modulin Derivatives. Journal of Biological Chemistry. 2011;286: 8933–8940. doi:10.1074/jbc.M111.221382

18. Zaman M, Andreasen M. Cross-talk between individual phenol-soluble modulins in Staphylococcus aureus biofilm enables rapid and efficient amyloid formation. Elife. 2020;9. doi:10.7554/eLife.59776

19. Marinelli P, Pallares I, Navarro S, Ventura S. Dissecting the contribution of Staphylococcus aureus α-phenol-soluble modulins to biofilm amyloid structure. Sci Rep. 2016;6: 1–13. doi:10.1038/srep34552

20. Tayeb-Fligelman E, Salinas N, Tabachnikov O, Landau M. Staphylococcus aureus PSMα3 Cross-α Fibril Polymorphism and Determinants of Cytotoxicity. Structure. 2020;28: 301–313.e6. doi:10.1016/J.STR.2019.12.006

21. Qi R, Joo HS, Sharma-Kuinkel B, Berlon NR, Park L, Fu C lung, et al. Increased in vitro phenol-soluble modulin production is associated with soft tissue infection source in clinical isolates of methicillin-susceptible Staphylococcus aureus. Journal of Infection. 2016;72: 302–308. doi:10.1016/j.jinf.2015.11.002

22. Berlon NR, Qi R, Sharma-Kuinkel BK, Joo HS, Park LP, George D, et al. Clinical MRSA isolates from skin and soft tissue infections show increased in vitro production of phenol soluble modulins. Journal of Infection. 2015;71: 447–457. doi:10.1016/j.jinf.2015.06.005

23. Yu J, Li M, Wang J, Hamushan M, Jiang F, Wang B, et al. Identification of Staphylococcus aureus virulence-modulating RNA from transcriptomics data with machine learning. Virulence. 2023;14. doi:10.1080/21505594.2023.2228657

24. Choe D, Szubin R, Dahesh S, Cho S, Nizet V, Palsson B, et al. Genome-scale analysis of Methicillin-resistant Staphylococcus aureus USA300 reveals a tradeoff between pathogenesis and drug resistance. doi:10.1038/s41598-018-20661-1

25. Briaud P, Zapf RL, Mayher AD, McReynolds AKG, Frey A, Sudnick EG, et al. The Small RNA Teg41 Is a Pleiotropic Regulator of Virulence in Staphylococcus aureus. Infect Immun. 2022;19. doi:10.1128/IAI.00236-22

26. Roberts C, Anderson KL, Murphy E, Projan SJ, Mounts W, Hurlburt B, et al. Characterizing the effect of the Staphylococcus aureus virulence factor regulator, SarA, on log-phase mRNA half-lives. J Bacteriol. 2006;188: 2593–2603. doi:10.1128/JB.188.7.2593-2603.2006

27. Lorenz R, Bernhart SH, Höner zu Siederdissen C, Tafer H, Flamm C, Stadler PF, et al. ViennaRNA Package 2.0. Algorithms for Molecular Biology. 2011;6: 26. doi:10.1186/1748-7188-6-26

28. Marinus T, Fessler AB, Ogle CA, Incarnato D. A novel SHAPE reagent enables the analysis of RNA structure in living cells with unprecedented accuracy. Nucleic Acids Res. 2021;49: e34–e34. doi:10.1093/nar/gkaa1255

29. Incarnato D, Morandi E, Simon LM, Oliviero S. RNA framework: An all-in-one toolkit for the analysis of RNA structures and post-transcriptional modifications. Nucleic Acids Res. 2018;46. doi:10.1093/nar/gky486

30. Wang R, Braughton KR, Kretschmer D, Bach THL, Queck SY, Li M, et al. Identification of novel cytolytic peptides as key virulence determinants for community-associated MRSA. Nature Medicine 2007 13:12. 2007;13: 1510–1514. doi:10.1038/nm1656

31. Cheung GYC, Joo HS, Chatterjee SS, Otto M. Phenol-soluble modulins - critical determinants of staphylococcal virulence. FEMS Microbiol Rev. 2014;38: 698–719. doi:10.1111/1574-6976.12057

32. Peschel A, Otto M. Phenol-soluble modulins and staphylococcal infection. Nat Rev Microbiol. 2013;11: 814. doi:10.1038/nrmicro3151

33. Schoenfelder SMK, Lange C, Prakash SA, Marincola G, Lerch MF, Wencker FDR, et al. The small non-coding RNA RsaE influences extracellular matrix composition in staphylococcus epidermidis biofilm communities. Otto M, editor. PLoS Pathog. 2019;15: e1007618. doi:10.1371/journal.ppat.1007618

34. Buzzo JR, Devaraj A, Gloag ES, Jurcisek JA, Robledo-Avila F, Kesler T, et al. Z-form extracellular DNA is a structural component of the bacterial biofilm matrix. Cell. 2021;184: 5740–5758.e17. doi:10.1016/J.CELL.2021.10.010

35. Stringer C, Wang T, Michaelos M, Pachitariu M. Cellpose: a generalist algorithm for cellular segmentation. Nature Methods 2020 18:1. 2020;18: 100–106. doi:10.1038/s41592-020-01018-x

36. Kaplan RM, Chambers DA, Glasgow RE. Big Data and Large Sample Size: A Cautionary Note on the Potential for Bias. Clin Transl Sci. 2014;7: 342–346. doi:10.1111/CTS.12178

37. Maher JM, Markey JC, Ebert-May D. The other half of the story: Effect size analysis in quantitative research. CBE Life Sci Educ. 2013;12: 345–351. doi:10.1187

38. O’Neill E, Pozzi C, Houston P, Smyth D, Humphreys H, Robinson DA, et al. Association between methicillin susceptibility and biofilm regulation in Staphylococcus aureus isolates from device-related infections. J Clin Microbiol. 2007;45. doi:10.1128/JCM.02280-06

39. Periasamy S, Joo HS, Duong AC, Bach THL, Tan VY, Chatterjee SS, et al. How Staphylococcus aureus biofilms develop their characteristic structure. Proc Natl Acad Sci U S A. 2012;109: 1281–1286. doi:10.1073

40. Mediati DG, Wong J, Gao W, McKellar SW, Pang I, Wu S, et al. RNase III-CLASH of multi-drug resistant Staphylococcus aureus reveals a regulatory mRNA 3’UTR required for intermediate vancomycin resistance. Nat Commun. 2022;13. doi:10.1038/s41467-022-31177-8

41. Busch A, Richter AS, Backofen R. IntaRNA: Efficient prediction of bacterial sRNA targets incorporating target site accessibility and seed regions. Bioinformatics. 2008;24: 2849–2856. doi:10.1093/bioinformatics/btn544

42. Tjaden B. TargetRNA3: predicting prokaryotic RNA regulatory targets with machine learning. Genome Biol. 2023;24: 276-. doi:10.1186/S13059-023-03117-2/FIGURES/5

43. Dugar G. Comprehensive architecture of the bacterial RNA interactome. bioRxiv. 2025; 2025.09.11.675593. doi:10.1101/2025.09.11.675593

44. Somerville GA, Chaussee MS, Morgan CI, Fitzgerald ‡ J Ross, Dorward DW, Reitzer LJ, et al. Staphylococcus aureus Aconitase Inactivation Unexpectedly Inhibits Post-Exponential-Phase Growth and Enhances Stationary-Phase Survival. Infect Immun. 2002;70: 6373–6382. doi:10.1128/IAI.70.11.6373-6382.2002

45. Zhu Y, Xiong YQ, Sadykov MR, Fey PD, Lei MG, Lee CY, et al. Tricarboxylic acid cycle-dependent attenuation of Staphylococcus aureus in vivo virulence by selective inhibition of amino acid transport. Infect Immun. 2009;77: 4256–4264. doi:10.1128/IAI.00195-09

46. Stevens E, Laabei M, Gardner S, Somerville GA, Massey RC. Cytolytic toxin production by Staphylococcus aureus is dependent upon the activity of the protoheme IX farnesyltransferase. Scientific Reports 2017 7:1. 2017;7: 13744-. doi:10.1038/s41598-017-14110-8

47. Soutourina O, Liu S, Lee AH, Caldelari I, Barrientos L, Mercier N, et al. Assembling the Current Pieces: The Puzzle of RNA-Mediated Regulation in Staphylococcus aureus. 2021. doi:10.3389/fmicb.2021.706690

48. Kreiswirth BN, Löfdahl S, Betley MJ, O’reilly M, Schlievert PM, Bergdoll MS, et al. The toxic shock syndrome exotoxin structural gene is not detectably transmitted by a prophage. Nature 1983 305:5936. 1983;305: 709–712. doi:10.1038/305709a0

49. Novick R. Properties of a cryptic high-frequency transducing phage in Staphylococcus aureus. Virology. 1967;33: 155–166. doi:10.1016/0042-6822(67)90105-5

50. Herbert S, Ziebandt AK, Ohlsen K, Schäfer T, Hecker M, Albrecht D, et al. Repair of global regulators in Staphylococcus aureus 8325 and comparative analysis with other clinical isolates. Infect Immun. 2010;78: 2877–2889. doi:10.1128/IAI.00088-10

51. Le Lam TN, Morvan C, Liu W, Bohn C, Jaszczyszyn Y, Bouloc P. Finding sRNA-associated phenotypes by competition assays: An example with Staphylococcus aureus. Methods. 2017;117: 21–27. doi:10.1016/J.YMETH.2016.11.018

52. McDougal LK, Steward CD, Killgore GE, Chaitram JM, McAllister SK, Tenover FC. Pulsed-Field Gel Electrophoresis Typing of Oxacillin-Resistant Staphylococcus aureus Isolates from the United States: Establishing a National Database. J Clin Microbiol. 2003;41: 5113–5120. doi:10.1128/JCM.41.11.5113-5120.2003

53. Schuster CF, Howard SA, Gründling A. Use of the counter selectable marker pheS* for genome engineering in staphylococcus aureus. Microbiology (United Kingdom). 2019;165: 572–584. doi:10.1099/mic.0.000791

54. Ivain L, Bordeau V, Eyraud A, Hallier M, Dreano S, Tattevin P, et al. An in vivo reporter assay for sRNA-directed gene control in Gram-positive bacteria: Identifying a novel sRNA target in Staphylococcus aureus. Nucleic Acids Res. 2017;45: 4994–5007. doi:10.1093/nar/gkx190

55. Geissmann Q. OpenCFU, a New Free and Open-Source Software to Count Cell Colonies and Other Circular Objects. PLoS One. 2013;8. doi:10.1371/journal.pone.0054072

56. Siegfried NA, Busan S, Rice GM, Nelson JAE, Weeks KM. RNA motif discovery by SHAPE and mutational profiling (SHAPE-MaP). Nat Methods. 2014;11: 959–965. doi:10.1038/nmeth.3029

57. Langmead B, Salzberg SL. Fast gapped-read alignment with Bowtie 2. Nat Methods. 2012;9: 357–359. doi:10.1038/nmeth.1923

58. Webb S, Hector RD, Kudla G, Granneman S. PAR-CLIP data indicate that Nrd1-Nab3-dependent transcription termination regulates expression of hundreds of protein coding genes in yeast. Genome Biol. 2014;15: R8. doi:10.1186/gb-2014-15-1-r8

59. Love MI, Huber W, Anders S. Moderated estimation of fold change and dispersion for RNA-seq data with DESeq2. Genome Biol. 2014;15: 550. doi:10.1186/s13059-014-0550-8

